# Environmental Drivers and Distribution of Pathogenic *Vibrio* Species in the Teign Estuary, UK

**DOI:** 10.64898/2026.06.30.735665

**Authors:** Hannah Boote, Nicola Coyle, Aoife Forde, Ioana Alexa, Molly Burchell, Stuart Reynolds, David J. Studholme, Sariqa Wagley

**Author notes:** Equal first author (Hannah Boote, Nicola Coyle).

## Abstract

Climate-driven increases in sea surface temperature have been associated with the expansion of *Vibrio* species and a corresponding rise in vibriosis cases in both human populations and aquaculture systems. Coastal waters across the south of England are increasingly becoming suitable for the growth and establishment of both human- and aquaculture-associated *Vibrio* species, potentially increasing vulnerability to the types of infections and disease outbreaks already reported elsewhere in the world. In this study, we report the presence of a diverse and well-established *Vibrio* community within the Teign Estuary, (Southwest, UK), including the human-pathogenic species *V. parahaemolyticus, V. cholerae* (non-O1/non-O139), *V. alginolyticus*, and *V. diabolicus*, as well as the important aquaculture pathogens *V. jasicida, V. aestuarianus*, and *V. anguillarum*. We identified *V. diabolicus*, a species that was indistinguishable from *V. alginolyticus* using conventional biochemical identification methods and could only be accurately resolved by whole-genome sequencing and developed novel PCR targets to differentiate these species in the lab. Using the insect infection model *Galleria mellonella*, we demonstrate that environmental isolates of *V. cholerae* (non-O1/non-O139), *V. parahaemolyticus*, and *V. alginolyticus* possess virulence potential. We also investigated the effects of sewage effluent on the growth of *Vibrio* isolates from the Teign Estuary and found that sewage can preferentially promote the growth of *Vibrio* species. Furthermore, several *Vibrio* isolates were multidrug resistant and carried antimicrobial resistance genes, highlighting the potential role of environmental *Vibrio* populations in the Teign Estuary as reservoirs of antimicrobial resistance. Together, these findings demonstrate how rising sea surface temperatures and sewage pollution may influence the emergence, persistence, and public health and aquaculture significance of *Vibrio* species in UK coastal waters.

## Introduction

Vibriosis is a gastrointestinal disease in humans caused by waterborne *Vibrio* species that can be fatal. It is also responsible for mass mortality events in populations of both wild and farmed shellfish worldwide. In humans, *V. parahaemolyticus* is the leading global cause of seafood-associated gastroenteritis, accounting for an estimated half a million cases annually [1]. Additionally, *V. cholerae* causes between 1.3 and 4.0 million cholera cases and an estimated 21,000–143,000 deaths each year.

In the United Kingdom, reported human infections caused by *Vibrio* species remain relatively low; the total number of annual confirmed laboratory reports were 141, 153 and 217 in the years 2022, 2023 and 2024 respectively [2]. Generally, these cases are travel-associated, while the true incidence of domestically acquired *Vibrio* infection is still uncertain. At present, around 22 *V. parahaemolyticus* infections are reported annually to the UK Health Security Agency (UKHSA), the majority of which are related to travel [3].

A recent study by the UKHSA analysed *V. cholerae* cases reported between 2004 and 2024 [4]. The authors identified 984 notifications over this period, corresponding to an average of approximately 51 cases per year. The highest annual number of cases occurred in 2010 (n = 74), while the lowest were recorded during the COVID-19 travel restrictions (8 cases in 2020 and 4 cases in 2021). Overall, 92.9% of cases were associated with international travel rather than domestic acquisition. The study also reported a small number of cases without a history of recent travel. In total, 14 cases were classified as not associated with foreign travel, while a further 56 cases had unknown travel histories. Specifically, three cases were identified during the COVID-19 travel restrictions that had no recorded travel outside of the UK [4]. These findings, along with the recent death of an 80-year-old man in Warwickshire with a non- O1/non-O139 *V. cholerae* infection (also not travel associated) suggest that a small proportion of *V. cholerae* infections in the UK may be domestically acquired, although the evidence remains limited.

In aquaculture settings, vibriosis has resulted in substantial production losses affecting shrimp, finfish, and shellfish industries, as well as large-scale mortality events in both managed and wild shellfisheries in countries including France, the USA, New Zealand, and Australia [5–10].

*Vibrio* species are predominantly found above 18°C, at temperatures that are suitable for their growth and proliferation [11, 12]. In a previous study, we analysed sea-surface temperature (SST) data from the global Operational Sea-Surface Temperature and Sea Ice Analysis (OSTIA) system to investigate temperature patterns around the coasts of England and Wales between 2015 and 2018 [13]. Among the locations examined was Teignmouth, Devon, UK. Our analysis showed that SSTs at Teignmouth remained below 18°C between 2015 and 2017 and fluctuated around 18°C during July and August 2018 [13]. Following publication of these findings, local fishermen operating within the River Teign reported observing summer water temperatures exceeding 19–20°C and expressed concerns that warming conditions may be creating a more favourable environment for *Vibrio* species (anecdotal evidence).

The Teign Estuary supports a range of recreational activities, including fishing, swimming, and water sports [14, 15]. Given its importance for public use, and the anecdotal evidence of warming temperatures, further investigation into temperature dynamics within the estuary and their potential influence on *Vibrio* ecology was carried out. The findings presented here demonstrate that pathogenic *Vibrio* species with the potential to cause disease are present at a low abundance within the Teign Estuary and the continued increase in SSTs in nearshore ecosystems along the south coast of England are suitable to support their growth and persistence.

## Results

### The Teign Estuary is warm enough to support growth of pathogenic *Vibrio* species

SST data was analysed at four coastal locations with proximity to Teignmouth (Weymouth, West Bay, Dawlish and Torbay), and Figure 1B shows there is an overall positive trend in number of days above 18°C from 2011 to 2025 around the southwest coast of England (Figure 1B). Our sample site in the Shaldon area of the Teign Estuary (in Teignmouth) is shown in Figure 1A. We collected SST data from a HOBO logger (Tempcon Instrumentation Ltd) at sampling location B (Figure 1A) between June 2024 and Oct 2025 (Figure 1C). In 2024, the average daily SST exceeded 18°C for a total of 23 days, in the months of July (4 days), August (15 days) and September (4 days). In 2025, the SST exceeded 18°C for a total of 57 days: 5 days in June, 16 days in July, the entire month of August (except for the 5^th^) and 6 days in September. Additionally, 21 days of temperature data were lost (logger error), from 22^nd^ June to 12^th^ July 2025. Therefore, it is likely that there were more than 57 days with a daily average >18°C, including days in this 21-day unrecorded period. Compared with 2024, there was a 2.4-fold increase in the number of days during which temperatures exceeded 18°C in 2025. This is consistent with 2024 being recorded as the coolest UK summer since 2015, and 2025 being the warmest UK summer on record [16]. For comparison, on the 21^st^ June (summer solstice) in 2024, the average temperature in the Teign Estuary was 16.16°C, whereas on the same date in 2025 the average was 19.19°C (+3.03°C difference). Daily maximum sea surface temperatures (SSTs) shown in Figure 1D indicate that, during the summer months, temperatures frequently exceeded 22°C for several hours each day. In 2025, daily maximum SSTs surpassed 25°C on four separate days. Laboratory studies have shown that *Vibrio* growth rate increases rapidly with temperature, while in the natural environment, the *Vibrio* abundance changes once temperatures pass certain critical levels [17]. These transient periods of elevated temperature may create localised thermal hotspots that promote the growth and persistence of *Vibrio* species within the estuary. Thus, short-term temperature maxima, rather than seasonal average temperatures alone, may play an important role in determining *Vibrio* abundance and the associated public health risk.

**Figure 1:**
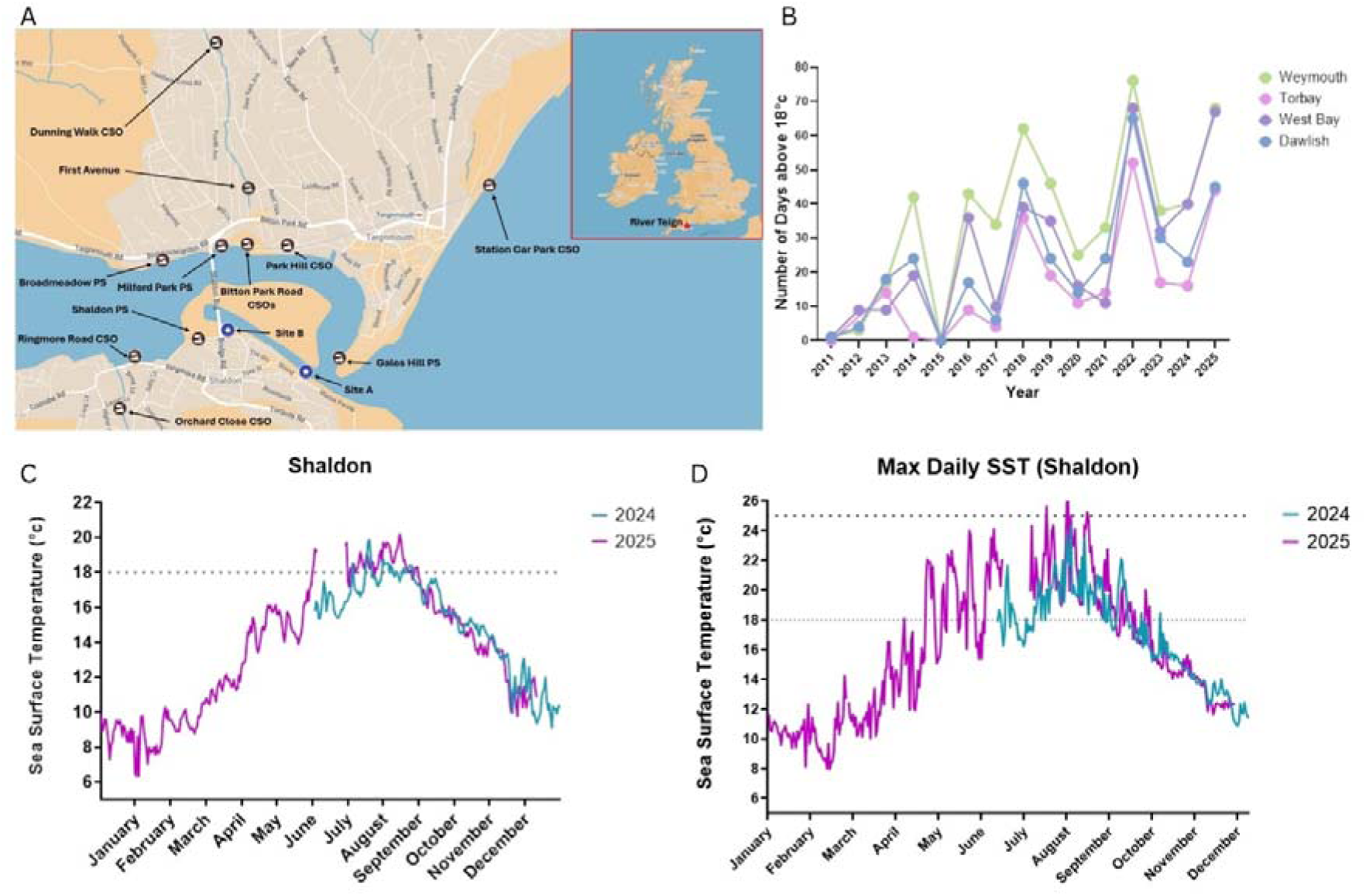
Sampling site in the Teign Estuary. **A**: Map of Teign Estuary, including Site A (EA/FORT sampling site) and Site B (our sampling site and where HOBO logger is situated) and sewage outflow pipes in brown symbols. **B:** The number of days where average sea surface temperature (SST) was greater than 18°C at nearby coastal locations across the Southwest including Weymouth, Torbay, West Bay and Dawlish. Data taken from the Channel Coastal Observatory, from 2011 to 2025. *[18]* **C:** Time series of daily SST data in the Teign Estuary. **D**: Time series of daily maximum SST data in the Teign Estuary. Blue and purple lines show SST in °C for 2024 and 2025, respectively. The data was from a HOBO logger (Tempcon Instrumentation Ltd).

### Water quality in the Teign Estuary is variable

Bathing water quality in England is assessed using concentrations of *Escherichia coli* (*E. coli*) and intestinal enterococci (IE), which are indicators of faecal matter. For coastal bathing waters to achieve an ‘Excellent’ classification, the 95^th^ percentile values must not exceed 250 CFU/100 mL for *E. coli* and 100 CFU/100 mL for IE. The official bathing water season in England runs from 15^th^ May to 30^th^ September, during which the Environment Agency (EA) undertakes routine monitoring [19, 20] [21]

In this study, we monitored coliform numbers for water (Figure 2A), sediment (Figure 2B) and shellfish (Figure 2C) alongside our monthly sampling from Site B (Figure 1A). We observed low coliform counts in all samples during the bathing season (May-September) increasing in number during the non-bathing season (October-April). Overall, the coliform counts were lowest in water, compared with sediment and shellfish. Coliform counts were highest within the shellfish tested, which is not surprising, as they are bivalve filter feeders and can bioaccumulate pathogens in their tissues and digestive tracts [22–24].

**Figure 2:**
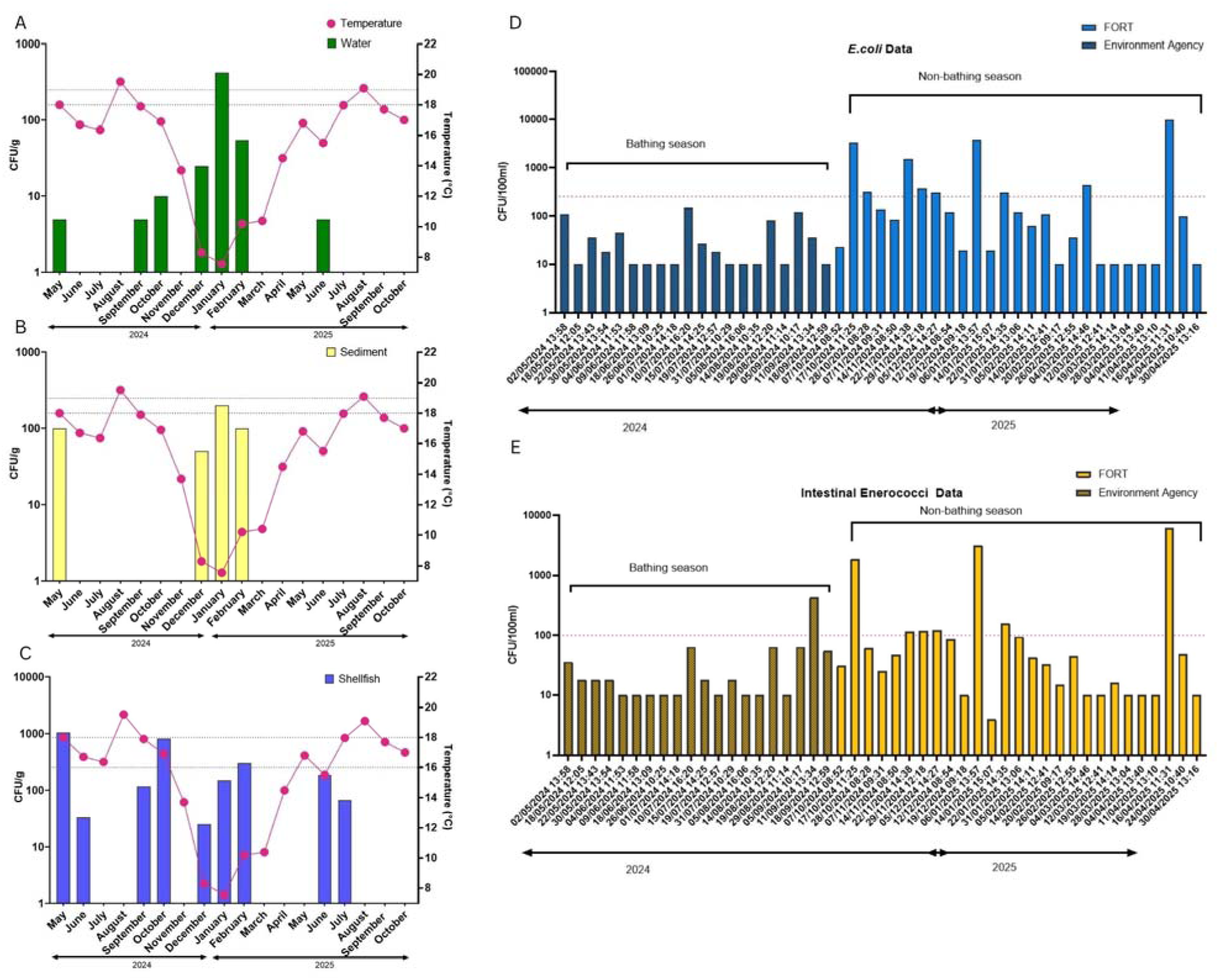
Enumeration of coliform bacteria in water. (A), sediment (B) and shellfish (C), and *Escherichia coli* (D) and intestinal enterococci (E) in water.

**Figure 3:**
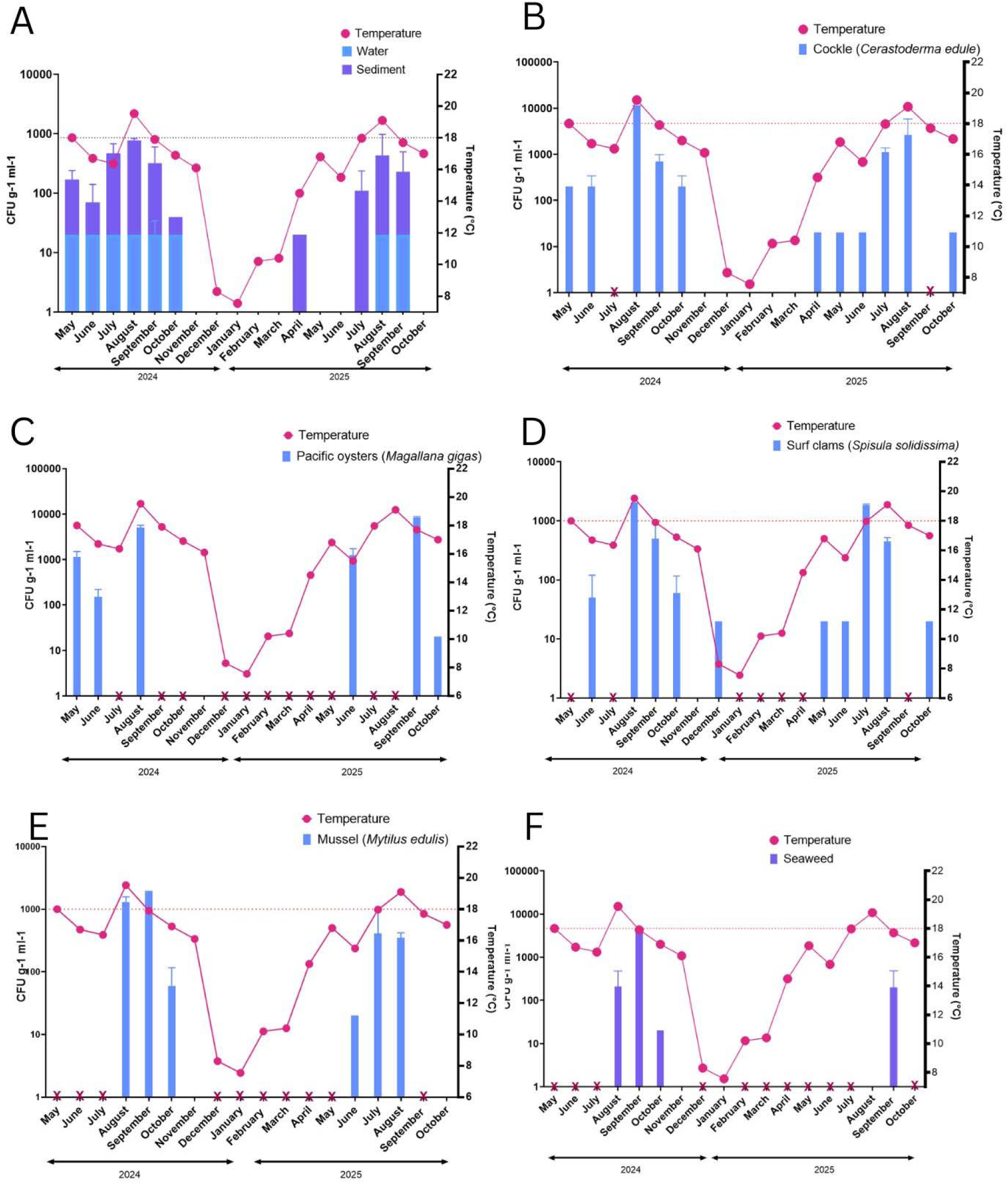
Abundances of *Vibrio* species in the Teign Estuary correlate with temperature. Bacterial abundance (CFU g^-1^) is plotted against temperature taken at the time of sampling. **A**: Water and sediment samples. **B** cockle. **C**: oyster. **D**: clam. **E**. mussel. **F** seaweed. The 18°C temperature is marked up by a red dotted line. A red bold X indicates when no sample was collected.

Additional data for water quality was provided by the local Friends of the River Teign (FORT) group, who collected water samples on randomised days every week at Site A (Figure 1A) in the Teign Estuary and sent them for commercial testing (Eurofins). This is the exact location from which the Environment Agency (EA) collect their water samples during the bathing season. According to both publicly available EA testing data and commercial testing from FORT, during the 2024 bathing season, *E. coli* numbers were below 200 CFU/100ml while IE numbers were also below 100 CFU/100ml except for two dates; September 2024 that showed an IE value at 460 CFU/100ml and on 31 July 2025 that showed an value of 1100 CFU/100ml *E Coli* and 1700 CFU/100ml IE (Figure 2D-E). Thus, data recovered from the EA and FORT data (Figure 2D-E) agreed with observations within this study that higher abundance of *E. coli* and IE can be seen during the non-bathing season (Oct-April), than in the bathing season. Higher numbers of coliforms are detected during the winter season compared to the summer season because of heavy rainfall. The heavy rainfall increases agricultural and urban run-off into waterways, leads to tidal mixing which brings in bacteria from the sediment into the water column and can trigger storm overflow events when sewage networks are overwhelmed [20, 25, 26].

### Culturable *Vibrio* species are present in the Teign Estuary between May and October

In total, 91 water, sediment and shellfish samples were collected in this study. Four initial samples were taken in 2023, and 41 and 46 samples were recovered in 2024 and 2025, respectively. Table 1 shows a summary of the samples tested in this study, including the lowest to highest number of *Vibrio* found in each sample. Further information on each of these samples has been detailed in Supplementary Table S1. The total *Vibrio* abundance was enumerated from plate counts, considering sucrose-utilising (yellow) and sucrose non-utilising (green) colonies on thiosulphate citrate bile sucrose agar (TCBS) plates as presumptive *Vibrio* colonies [27]. The total abundance of presumptive *Vibrio* colonies in all the water samples tested, was low, ranging between 0 and 20 CFU ml^-1^, regardless of temperature. The total abundance of *Vibrio* in the sediment samples was higher than in the water samples and peaked in August 2024 (770 CFU g^-1^) and August 2025 (430 CFU g^-1^), which coincided with peaks in temperature. Shellfish samples showed the highest burden of *Vibrio*, with colony counts peaking when temperatures were warmest in the Teign Estuary. In 2024, the highest abundance levels of *Vibrio* CFUs were seen in a cockle sample (10950 CFU g^-1^) and in 2025, the highest colony count was observed in an oyster sample tested in September (8400 CFU g^-1^).

**Table 1:** Summary of water, sediment, shellfish and seaweed samples tested in this study.

| Type of sample | Water | Sediment | *Cerastoderma edule*<br>(Common cockle) | *Magallana gigas*<br>(Pacific Oyster) | *Spisula solidissima*<br>(Surf clam) | *Mytilus edulis*<br>(Blue mussels) | Seaweed |
| --- | --- | --- | --- | --- | --- | --- | --- |
| Number of samples tested | 20 | 19 | 17 | 8 | 11 | 7 | 8 |
| Range of vibrios from lowest to highest number found.<br>CFU g<sup>-1</sup> ml<sup>-1</sup> | 0 - 30 | 20 - 400 high in Aug 2024 and Aug 2025 | 20 - 12350 high in Aug 2024 | 20 - 8400 high in Aug 2024 and Sept 2025 | 20 - 3200 high in July 2025 | 20 - 2900 high in Aug 2025 | 200 - 4100 high in Sep 2024 |
| *V. parahaemolyticus* | + | + | + | + | + | + | + |
| *V. cholerae* | + | - | - | - | + | - | - |
| *V. alginolyticus* | + | + | + | + | + | + | + |
| *V. diabolicus* | + | + | + | + | + | - | + |
| *V. jasicida* | + | + | + | - | + | - | - |
| *V. aestuarianus* | - | + | + | - | - | - | - |
| *V. neptunius* | + | - | - | - | - | - | - |
| *V. anguillarum* | + | - | - | - | - | - | - |
| *V. alfacensis* | - | - | + | - | - | - | - |
| *V. pacinii* | - | - | + | - | - | - | - |
| *Vibrio* species | - | - | - | - | + | - | - |

We performed whole-genome sequencing on 53 presumptive *Vibrio* isolates and classified them to species level with high confidence. A summary of the *Vibrio* species isolated and identified from at least one sample can be found in Table 1. Included in these isolates was a *V. alginolyticus* strain isolated from a patient with a wound infection in Guernsey in the Channel Islands, British Isles in 2011 which has not been sequenced before [28].

### Detection of pathogenic species *V. parahaemolyticus* and *V. cholerae* non-O1/O139 in Teign Estuary

Twelve *Vibrio* species are known to cause disease in humans, including *V. parahaemolyticus*, *V. cholerae* and *V. alginolyticus* [29]. We isolated *V. parahaemolyticus* at low abundances in a range of samples throughout 2024, and 2025. In 2024, we measured *V. parahaemolyticus* in August, September and October, ranging in values from <20 – 1000 CFU g^-1^ ml^-1^ in 10 samples (water, sediment, cockles, surf clams, blue mussel and seaweed). In 2025, *V. parahaemolyticus* was detected in June through to October at levels of less than 100 CFU g^-1^ ml^-1^ in nine samples (sediment, Pacific oysters, blue mussels, cockles, and surf clams).

All putative isolates of *V. parahaemolyticus* were confirmed by PCR, using the species marker *toxR* [30]. In addition, the presence of the virulence genes *tdh* and *trh* was determined by PCR and found to be negative [31]. Analysis of genome sequences for 12 of these isolates showed that they were representatives of four different multi-locus sequence-types, each falling into a different phylogenomic clade within the wider *V. parahaemolyticus* population (Figure 4, Supplementary Figure S2). No clear relationship was observed between sequence type and either the year of isolation nor the source of isolation. For example, sequence type 1366 was represented by six isolates, recovered in both 2024 and 2025 and from mussels, sediment, and surf clams. These findings suggest that genetically diverse *V. parahaemolyticus* strains are present in the Teign Estuary and that some sequence-types persist and reoccur across multiple years. While the three of the four lineages of *V. parahaemolyticus* were distinct within the species population, one isolate, EXE 26/2024-C, was highly similar to an isolate from the UK. This isolate was submitted by the Gastrointestinal Bacteria Reference Unit at UKHSA, isolated from a food source in August 2024, as part of their routine sequencing of *Vibrio* species (GCA_041560915.1) (Supplementary Figure S1).

**Figure 4.**
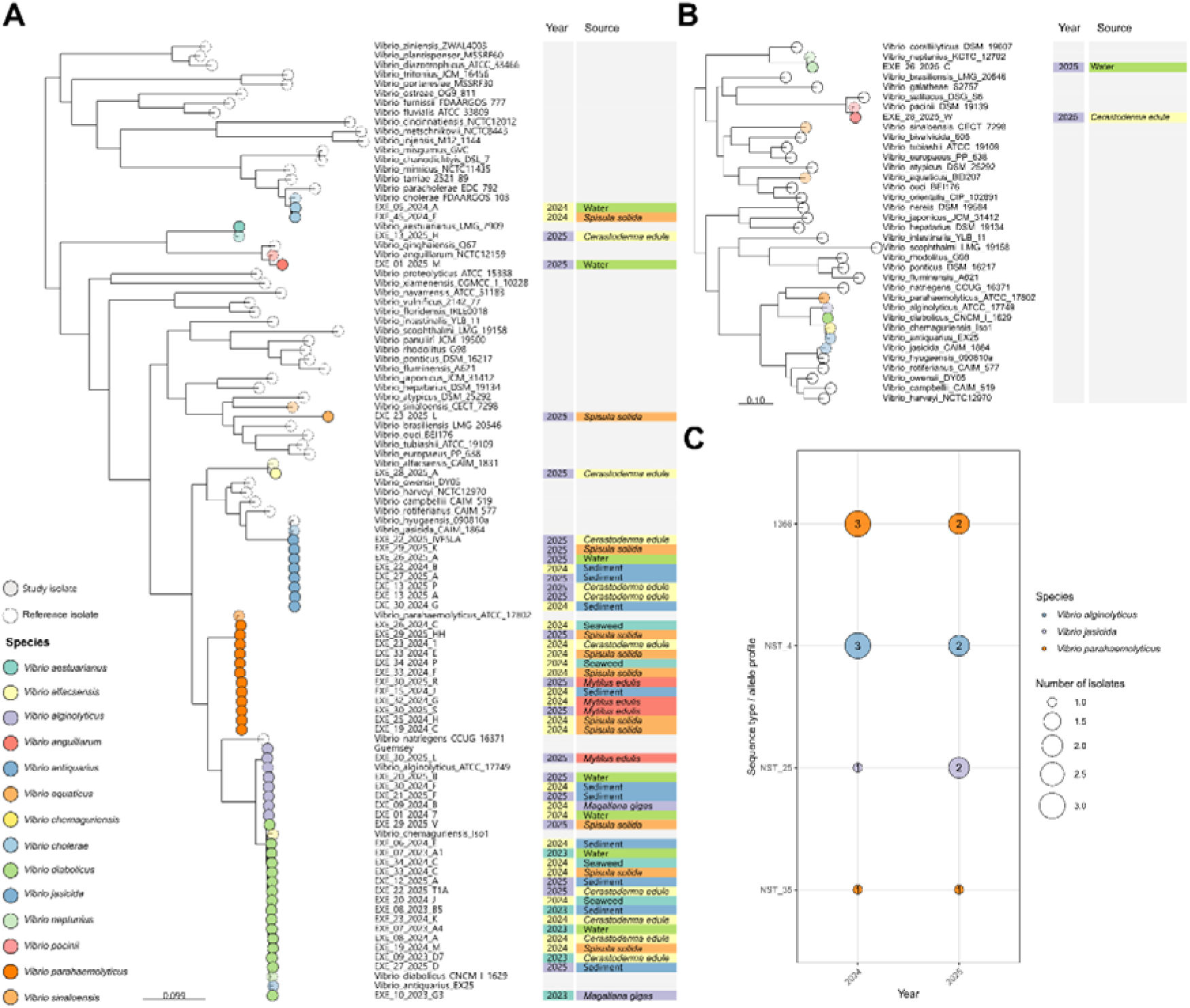
Phylogeny of 53 study isolates and representative isolates of species within the *Vibrio* genus produced by Phame. Two phylogenies were constructed using PhaME to help confirm species identities of each whole genome sequenced in this study. **A**: Tree including most isolates in the study. **B**: Tree including study isolates from species *V. neptunius* and *V. pacinii*. **C**: Number of isolates from multi-locus sequence types (STs) that are shared across years 2024. STs with allelic profiles not found in the MLST schemes used are denoted as new STs (NST_XX).

We also detected *V. cholerae* at low levels (<20 CFU ml^-1^) in June 2024 (from a water and cockle sample) and in December 2024 (from surf clams). The strains were identified as *V. cholerae* by the API20E profile and further confirmed by WGS (Figure 4). Strains of *V. cholerae* belonging to serogroup O1 biotype El Tor and serogroup O139 have been described as causative agents of cholera [32–34]. To cause cholera diarrhoea, the bacteria require genes encoding several virulence factors including cholera toxin (*ctxA*), classical or El Tor colonisation toxin-coregulated pilus (*tcpA*), and virulence regulator (*toxR*) [35–38].

We scanned the sequences of the two *V. cholerae* isolates from the Teign Estuary for virulence genes, using ABRicate and BLASTN searches [39] [40]. Although both genomes contained *toxR*, neither genome contained *ctxA* nor *tcpA*. Both isolates lacked the *wbeO1* and *wbfO139* gene clusters and therefore can be categorised as non-O1 and non-O139. *V. cholerae* non-O1 and non-O139 strains can cause sporadic outbreaks of serious gastroenteritis but that is distinct from cholera [41] [42–45]. These two *V. cholerae* isolates from the Teign Estuary were genetically divergent, belonging to different multi-locus sequence types: EXE 05/2024-A isolated from water in June 2024 was identified as ST495 and was closely related to isolate VN10012 (GenBank: GCA_021014825.1), which was previously isolated from oysters in Germany in 2011. EXE 45/2024-E, isolated in December 2024 from surf clams, was ST466 and was closely related to two UK isolates 273589 and 529272 (from 2018) with genome sequences deposited by Public Health England (GenBank GCA_015812805.1 and GCA_015802055.1).

### *V. alginolyticus* and *V. diabolicus* are present in the Teign Estuary

*V. alginolyticus* and *V. diabolicus* are closely related species in the *V. harveyi* clade [46]. During our environmental sampling, we isolated bacteria that matched the phenotypic characteristics of *V. alginolyticus*, a species commonly found in marine and estuarine environments worldwide that is an important pathogen in aquaculture and an opportunistic pathogen of humans [47–52]. These swarming and sucrose-fermenting bacteria form large, yellow colonies on TCBS agar. Their metabolic profiles, identified using API, identified them as *V. alginolyticus* with a high degree of certainty. We recovered such isolates during the summer months in 55 samples from 2024 and 2025. Whole genome sequencing analysis of 24 of these isolates revealed that while six isolates belonged to *V. alginolyticus* as expected, 17 instead belonged to the closely related species *V. diabolicus*.

To further explore the species classifications of these isolates, we placed them in a phylogeny of publicly accessible isolates submitted to the NCBI as *V. alginolyticus*, *V. diabolicus*, or its synonym *V. antiquarius*. This confirmed the observed species classifications, and the diversity of strains isolated in the Teign Estuary became apparent, with three lineages of *V. alginolyticu*s and six distinct *V. diabolicus* lineages (Supplementary Figure S2). One lineage of *V. alginolyticus* containing five isolates (NST_4) was observed, appearing between May 2024 and June 2025 in a variety of sources, water (n=2), sediment (n=2) and oysters (n=1) (Figure 4C). Two isolates EXE 22/2025-T1A, assigned to ST31 using the *Vibrio* MLST scheme, and EXE 20/2024-J which differs from ST31 by one allele (*pyrH*), both closely resemble three *V. diabolicus* genomes including PHW7 (ST31), isolated in an estuary on the southwest coast of the UK in September 2020 (GCF_041894665.1), HS-43-7A (GCF_024748135.1) isolated from an oyster in Canada in 2016, and V2 (GCF_001010935.1) isolated in a Dentex dentex host experiencing vibriosis. Two isolates (EXE_33_2024_C and EXE_34_2024_C) assigned to ST32, were recovered in 2024 from seaweed and *Spisula solida,* respectively, and clustered with one publicly accessible genome (HS-40-3, GCF_024748335.1) found in 2015 from clams in Canada.

Given the close phylogenetic relationship and the phenotypic similarity between *V. diabolicus* and *V. alginolyticus* [46], it is not surprising that some genome sequences of these species have been misidentified in the public sequence repositories. In the GenBank database, on 20^th^ June 2026 we identified 401 genome assemblies that phylogenetically fall within *V. diabolicus [53]*. Of those, the majority (n=297) were correctly identified as *V. diabolicus*. The remaining genome assemblies were labelled as *V. alginolyticus* (n=14), *V. antiquarius* (n=44), *V. chamguriensis* (n=10), *V. parahaemolyticus* (n=2) and unidentified *V. species* (n=35). Conversely, only a single *V. alginolyticus* isolate appeared to be mis-attributed to *V. diabolicus* [46]. These findings indicate that taxonomic confusion is ongoing. Contributing to the confusion, 44 genome assemblies in the GenBank are denoted as “*V. antiquarius*” [54, 55]. Although it has been claimed that this species name is synonymous with *V. diabolicus* [46], according to the List of Prokaryotic names with Standing in Nomenclature (LPSN) [56] the name “*V. diabolicus*” is not validly published (and is therefore written in quotation marks). Ten genome assemblies are labelled in the public databases as “*Vibrio chemaguriensis*” [57]. The LPSN notes that this name is not validly published, despite being proposed in a previous publication[58]. Phylogenetically, these genomes fall within *V. diabolicus*.

We developed a PCR assay to distinguish the two closely related species *V. diabolicus* and *V. alginolyticus*, based on four protein-coding genes previously proposed as species markers for *V. diabolicus* [46]: hypothetical protein (ACY49842.1), hydroxyectoine utilisation dehydratase (ACY53324.1), ornithine cyclodeaminase (ACY53325.1) and phosphoesterase (ACY53763.1). PCR assays based on ACY53325.1 tested positive for all our *V. diabolicus* isolates (n=17) and PCR negative for all *V. alginolyticus* isolates (n=8). On the other hand, two assays yielded false positives and negatives. EXE 21/2025-F tested positive for ACY53324.1 and EXE 30/2024-F tested positive for ACY53763.1, while EXE 29/2025-V was negative for ACY53324.1. PCRs based on the gene ACY49842.1 could not identify *V. diabolicus* since all *V. diabolicus* and *V. alginolyticus* isolates were positive for this marker. This indicates that for distinguishing between *V. alginolyticus* and *V. diabolicus* in the lab, the most effective PCR assay is based on ACY53325.1 and using it in combination with secondary markers ACY53324.1 and ACY53763.1 may increase confidence in species assignments.

### Beyond human pathogens: whole genome sequencing reveals diverse *Vibrio* species in the Teign Estuary

Previous studies looking at the presence of *Vibrio* species in UK waterways and shellfish have focused on human pathogenic species, including *V. cholerae, V. parahaemolyticus* and *V. vulnificus.* This is primarily because methods for identifying these pathogens are well established alongside their public health risk. However, biochemical methods for rapid identification of non-human pathogenic *Vibrio* species remain inadequate. The use of API20 or 20NE strips for identifying and differentiating bacteria within the Enterobacteriaceae are constrained by the limitations of its reference database. In this study, we found colonies appearing on TCBS that were sucrose fermenters and non-fermenters but were not identifiable via the API system. We used whole genome sequencing to identify 15 unidentified species that were appearing in our samples. We identified eight of them as *V. jasicida*, and the others as *V. anguillarum, V. aestuarianus, V. neptunius, V. alfacsensis, V. pacinii* and an unidentified *Vibrio* species*. V. jasicida* was first described in 2012 when it was recovered from marine invertebrates and vertebrates, including Atlantic salmon and flounder [59]. *V. jasicida* previously classified as *V. harveyi* has caused mortality rates of >75% for phyllosoma larvae of the packhorse rock lobster *Jasus verreauxi* [60]. It has been isolated previously from European flat oysters (*Ostrea edulis*) and from Pacific oysters (*Magallana gigas*, formerly *Crassostrea gigas*) from the south coast of UK [13]. *V. anguillarum* is a well-established marine pathogen and can cause severe haemorrhagic septicaemia affecting a wide range of marine and estuarine fish species [5, 61–63]. *V. aestuarianus*, an important pathogen of oysters that has been linked to significant summer mortality events in cultivated and wild oyster populations in Ireland, France and Spain [64] [65–69]. *V. neptunius* has been isolated from numerous marine animals and is a pathogen of oysters and clams [70–72]. *V. alfacsensis* has been potentially identified as a threat to turbot (*Scophthalmus maximus*) and has a strong tendency to carry AMR genes [73]. *V. pacinii* has previously been isolated from healthy shrimp larvae (*Penaeus chinensis*) in Shandong Province, China, Atlantic salmon (*Salmo salar*) in Tasmania, sea bass (*Dicentrarchus labrax*) in Spain and from goldsinny wrasse (*Ctenolabrus rupestris*) [74, 75]. Along with these *Vibrio* species mentioned above, *V. alginolyticus* is an opportunistic marine pathogen that can cause septicaemia in fish and shellfish. This species has been associated with high-mortality outbreaks in aquaculture populations of fish, oysters, and shrimp[51, 76–78]. Collectively, these findings demonstrate that the *Vibrio* community present within the Teign Estuary extends beyond traditionally monitored human pathogens and includes several species associated with disease in marine animals. This highlights the need for broader surveillance approaches that consider both public health and marine ecosystem health.

### Pathogenic potential of V. parahaemolyticus, V. cholerae, V. alginolyticus and *V. diabolicus* strains from the Teign Estuary

Previous work has shown that the insect infection model *Galleria mellonella* can be used to assess the virulence of *V. parahaemolyticus* strain [79]. The *G. mellonella* model can distinguish between clinical toxigenic *V. parahaemolyticus* strains and non-toxigenic strains, which usually predominate in the environment. In this work, we tested 2 representative strains of *V. parahaemolyticus* (EXE 19/2024-C and EXE 24/25-E), *V. alginolyticus* (EXE 09/2024-B and EXE 30/2025-L), *V. diabolicus* (EXE 20/2024-J and EXE 27/2025 – D) and *V. cholerae* (EXE 05/2024-A, EXE 45/2024-E) in the *G. mellonella* model. As a comparison, we also injected *V. parahaemolyticus* strain G35, a well-referenced clinical TDH and TRH positive strain that has been shown to be highly virulent in the *G. mellonella* model.

Both *V. cholerae* strains displayed very high virulence in larvae model with no larvae surviving after 24 hours at very low doses (83-330 cells - see Supplementary Table S2) (Figure 5A). Both strains of *V. cholerae* identified from the Teign Estuary were non-O1/non-O139 serotype and do not encode the genes for cholera toxin that lead to the cholera infection that is caused by O1/O139 serotypes. Non-O1/non-O139 strains are often associated with sporadic but serious gastroenteritis cases [4, 42, 45], and these strains may harbour a range of alternative virulence factors, including haemolysin, repeats-in-toxin (RTX), heat-stable enterotoxin, haemagglutinin/protease, type III secretion systems (T3SS), and type VI secretion systems (T6SS). WGS of *V. cholerae* EXE 05/2024-A and EXE 45/2024E revealed the presence of key virulence factors that may be responsible for the pathogenic potential of these two strains observed in the *G. mellonella* model including RTX genes, T6SS genes, hemagglutinin/proteinase HapA and HlyA / *V. cholerae* cytolysin (VCC) ([80–83]. In *V. cholerae* EXE 05/2025A, an additional virulence gene encodes a cholix toxin (*chxA),* which has been shown to halt protein synthesis in eukaryotic cells [84, 85]. Collectively, the presence of these virulence genes in the *V. cholerae* isolates from the Teign Estuary could account for the virulence we observed in the larvae infection model.

**Figure 5:**
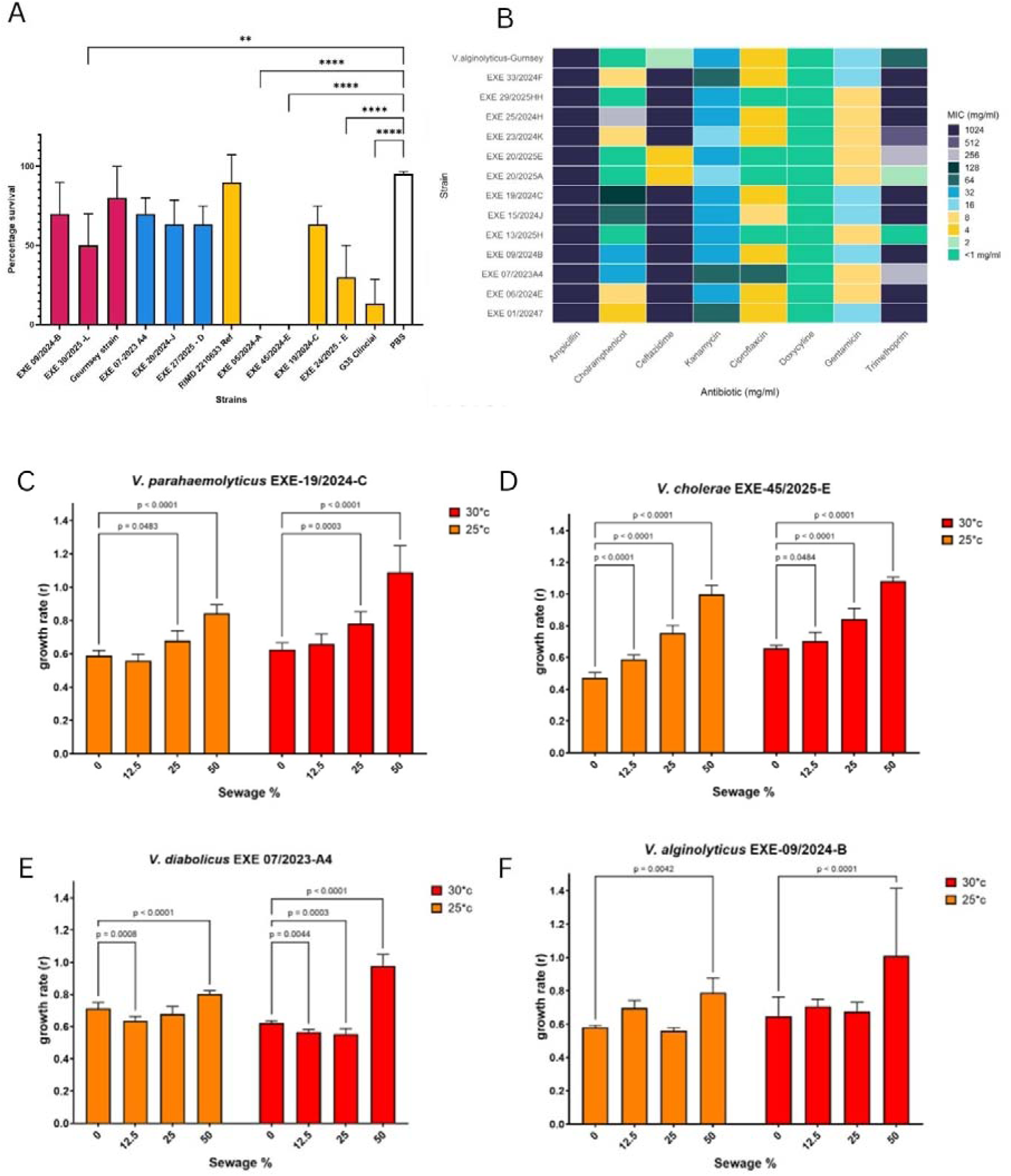
A: Survival of *Galleria mellonella* 48 h after challenge with ∼10^2^–10^3^ CFU of 12 *Vibrio* strains. *Vibrio* strains were grown at 37°C and percentage survival was measured after 48 h. The results shown here are the means of three experiments, each using groups of 10 larvae per strain. The error bars indicate standard deviation. B: Heatmap to depict the MIC results for 13 Vibrio strains isolated from the Teign Estuary. C-F: Growth rates (r) for four *Vibrio* species strains when co-cultured with raw sewage concentrations (50%, 25%, 12.5% and no sewage (0)). Strains were grown at 30 or 25°C for 12 hours.

The *V. parahaemolyticus* EXE 19/2024-C and EXE 24/2025E showed 60% survival and 30% survival, respectively (when injected with doses between 80-1435 cells) after 48 hrs (Figure 5A). Both strains were non-toxigenic, i.e. negative for haemolysins TDH and TRH that are well established as virulence factors in *V. parahaemolyticus [31, 86]*. One of the *V. alginolyticus* strains (EXE 30/2025-L) tested showed 50% survival after 48 hrs (when injected with a dose between 100-965 cells). Consequently, increases in the abundance of these strains in the Teign Estuary within the water and shellfish populations may elevate the risk of disease to human exposure and should be a public health concern. Taken together, the results demonstrate that the *V. cholerae*, *V. parahaemolyticus*, and *V. alginolyticus* isolates tested were highly virulent in the larval infection model, highlighting their potential pathogenicity.

### Co-culture with sewage can increase the growth of *Vibrio* species

The growth of four *Vibrio* species strains identified from the Teign Estuary were tested in the presence of filtered sewage at various concentrations and temperatures. Growth rates increased significantly in the presence of sewage for all four strains tested although the response was variable per strain (Figure 5C-F). The biggest impact was seen at 50% sewage concentration where a significant increase in growth rate (r) was observed compared to the no sewage control at both 25°C and 30°C. In the presence of 25% sewage a significant growth rate increase was observed in *V. parahaemolyticus* EXE 19/2024-C (Figure 5C) and in *V. cholerae* EXE- 45/2025-E (Figure 5D) at both 25°C and 30°C, while with *V. diabolicus* EXE 07/2023-A4 (Figure 5E) a significant increase was only seen at 30°C. In the presence of 12.5% sewage concentration, only *V. cholerae* EXE- 45/2025-E (Figure 5D) increased its growth rate compared to the no sewage control at both 25°C and 30°C.

To analyse differences between growth rates across conditions, a two-way ANOVA and Dunnett’s multiple comparisons test were performed (Supplementary Figure S3). Overall, the growth rate of the different *Vibrio* species tested was substantially influenced by temperature and the addition of sewage at different concentrations. (Supplementary Figure S3). For *V. cholerae* EXE 45/2024-E, the increases in growth rate observed were highly associated with the addition of sewage, with sewage concentration explaining 84% of variation seen. A further 9% of the variation could be explained by changes in temperature. Strains grown at 30°C showed consistently higher growth rates indicating a positive effect of temperature on these strains.

Similarly, for both *V. parahaemolyticus* EXE 19/2024-C *and V. diabolicus* EXE 07/2023-A4, a large portion of the variation seen was explained by sewage concentration (68 to 64.4% respectively). The growth of *V. parahaemolyticus* EXE-19/2024-C was most affected by temperature, accounting for 11.9% of the variation seen, likely due to the increased growth seen at 30°C particularly at 25% and 50% sewage concentrations.

While *V. alginolyticus* EXE 09/2024-B experienced a clear increase in growth rate at 50% sewage concentration at 30°C, this boost in growth was less pronounced at 25°C, and growth was not significantly different from the controls at lower sewage concentrations. These inconsistent changes in growth rate in response to sewage and temperature indicate that while these factors do influence growth rate, they explain a smaller portion of the variation seen in this strain.

Together these results indicate that sewage inputs and elevated temperature promote the growth of these four strains, with the most consistent positive impact seen in *V. cholerae* EXE 45/2024-E. For other strains, growth rate increases were most pronounced at 50% sewage concentration. Together these results indicate that sewage outflow events during the summer could lead to the proliferation of *Vibrio* species in the Teign Estuary.

### *Vibrio* strains show resistant profiles to multiple antibiotics

To determine whether *Vibrio* species isolated from the Teign Estuary were susceptible or resistant to antibiotics, we performed antimicrobial susceptibility testing using a 96-well plate microdilution technique against the following antibiotics: kanamycin, ampicillin, gentamicin, chloramphenicol, trimethoprim, ceftazidime, ciprofloxacin and doxycycline hyclate. We chose *V. alginolyticus* (n = 3)*, V. diabolicus* (n = 3), *V. parahaemolyticus* (n = 6) and *V. aestuarianus* (n = 1) strains isolated from the Teign Estuary between 2023 and 2025. In our strain panel, we included a clinical *V. alginolyticus* strain that was isolated from a patient in Guernsey with a wound infection [28]. The minimum inhibitory concentration (MIC) was defined as the lowest antibiotic concentration at which no visible growth occurred. Supplementary Table S3 shows the list of strains tested and the antibiotic MIC outcome, while Figure 5B shows these same results illustrated in a heatmap.

All strains were resistant to ampicillin at concentrations of 1024 mg/ml and sensitive to doxycycline at concentrations of <1mg/ml. Six of the seven *V. parahaemolyticus* isolates and two of the three *V. alginolyticus* tested showed resistance to 1024 mg/ml of trimethoprim. All isolates tested were resistant to ceftazidime except *V. alginolyticus* EXE 20/2025–A and *V. parahaemolyticus* EXE 20/2025 – E.

All strains tested had multiple antibiotic resistance (MAR) indices well above the critical threshold of 0.2 (Supplementary Table S3), with values ranging from 0.5 to 0.75. This indicates that the 13 *Vibrio* strains tested from the Teign Estuary are multidrug resistant and underlines the risk that environmental *Vibrio* species from the Teign Estuary pose as antimicrobial resistance (AMR) reservoirs. For the sequenced *Vibrio* isolates, we identified AMR genes in the genomes using AMRFinderPlus [87]. The *tet(35)* gene was present in nearly all isolates with AMR hits, whereas the β-lactamases were species-specific (e.g. CARB-42 in *V. alginolyticus*, *CARB-18/20/21* in *V. parahaemolyticus*, and CARB-57-like enzymes in *V. diabolicus*) see Table 2. The *tet(35)* gene was widely distributed among the isolates analysed and has previously been suggested to represent an intrinsic resistance determinant in several *Vibrio* species [88] [89] [90].

**Table 2:** Antimicrobial resistance determinants identified by AMRFinderPlus in *Vibrio* isolates. . Antimicrobial resistance (AMR) determinants identified in the genomes of the *Vibrio* isolates using AMRFinderPlus [87]. The table summarises the AMR genes detected within each species and their predicted antimicrobial classes. The number of isolates carrying each determinant is indicated where applicable. Genes associated with tetracycline resistance (tet(35)) were the most frequently detected, whereas β-lactamase genes showed species-specific distributions.

| Species | No.<br>isolates | AMR determinants detected | Predicted resistance |
| --- | --- | --- | --- |
| *Vibrio aesturianus* | 1 | None detected | — |
| *Vibrio alfacensis* | 1 | *tet*(35) | Tetracycline |
| *Vibrio alginolyticus* | 7 | *tet*(35), *bla*CARB-42 | Tetracycline; β-lactams |
| *Vibrio anguillarum* | 1 | None detected | — |
| *Vibrio cholerae* | 2 | *almE*, *almF*, *almG*; *varG* (1 isolate) | Colistin; carbapenem-associated β-lactam<br>resistance |
| *Vibrio diabolicus* | 13 | *tet*(35), *bla*CARB / *bla*CARB-57, *fos* (3 isolates), *qnrC* (1<br>isolate) | Tetracycline; β-lactams; fosfomycin; quinolones |
| *Vibrio jasicida* | 8 | *tet*(35); *bla*VHW (1 isolate) | Tetracycline; β-lactams |
| *Vibrio neptunius* | 1 | None detected | — |
| *Vibrio pacinii* | 1 | None detected | — |
| *Vibrio parahaemolyticus* | 11 | *tet*(35); *bla*CARB-18, *bla*CARB-20, *bla*CARB-21 | Tetracycline; β-lactams |
| *Vibrio* sp. | 1 | None detected | — |

## Discussion

### The River Teign and Estuary

The River Teign stretches approximately 50 kilometres, flowing from the uplands of Dartmoor to its mouth at Teignmouth on the south Devon coast. Winding through a steep-sided valley, the river traces a southward course along the edge of Dartmoor National Park before emptying into the Teign Estuary and eventually reaching the sea. The Teign Estuary is made up of a broad, east - west aligned tidal river channel. The Estuary is not currently designated as a Marine Protected Area (MPA). Shellfish harvesting and aquaculture have taken place in Devon’s estuaries for centuries, with mussels and oysters being the primary species harvested. In the Teign Estuary, commercial harvesting of mussels (*Mytilus edulis*) and Pacific oysters (*Magallana gigas*, formerly *Crassostrea gigas*) has been carried out until recently. Cockles (*Cerastoderma edule*) are also found in the estuary and have been collected at low levels historically and continue to be so today. However, unlike mussels and Pacific oysters, cockle populations in the Teign have never been abundant enough to support commercial harvesting. As well as shellfish, the River Teign and estuary have been used for commercial netting of salmon and sea trout. In September 2023, local fishermen shared their concerns with us (at the University of Exeter) about how nature is adapting around them in response to climate change. Noting that sea surface temperatures in the Teign Estuary were reaching 18.5°C as early as June whereas historically such temperatures were not typically observed until late September. They also described a range of other environmental changes they had witnessed over the years. The Teign Estuary was home to a significant salmon (*Salmo salar*) and sea trout (an anadromous form of *Salmo trutta*) fishery. Both salmon and sea trout (locally known as peal) were previously caught by seine nets towed by rowing boats, but now the net fishery is reduced and restricted to a limited season during which only sea trout may be retained. The Teign Estuary serves as an important nursery habitat for European bass (*Dicentrarchus labrax*), with juveniles remaining in the estuary for several years before migrating to deeper offshore waters. Adult bass also return during the summer months to exploit abundant prey resources. As a species sensitive to temperature, fisherman have given anecdotal reports of bass being caught later in the year which may reflect increasing estuarine water temperatures, although recreational angling remains permitted within the estuary. The Teign Estuary historically supported populations of European flounder (*Platichthys flesus*) fishery including the UK capture record in 1994. Once attracting anglers from across the UK, National Flounder Championships catches regularly exceeded several hundred fish; however, recent competitions have recorded substantially lower catches indicating a marked decline in flounder population in the estuary. The decline in the flounder population over the years is contrasted by a noticeable rise in gilthead bream (*Sparus Aurata*) which are regarded as a warm water species with the southern coast of the British Isles being previously seen as the limit of its range. However, the species is now a regular target for some recreational anglers. Fishermen also observe the appearance of the invasive species *Grateloupia turuturu* commonly called devil’s tongue weed, a marine species of Rhodophyta (red algae), which was first observed in 2021 and has since become more widespread and established in the river and estuary. Additionally, fishermen reported occasional increases in small bait fish appearing as early as June—an occurrence usually not seen until September. The anecdotal observations from local fishermen that nature was adapting around them could be attributed to the increase in sea surface temperatures in the River Teign and Estuary occurring in recent years.

In 2022, we published data from OSTIA system [1], demonstrating SST patterns in Teignmouth, Devon, UK remained below 18°C between 2015 and 2017. However, in 2018 temperatures fluctuated around 18 °C during July and August and concluded that conditions in the Teign Estuary were not suitable to support the growth of *Vibrio* species at that time [2]. Our current study shows the SST in the Teign Estuary now are readily above 18 °C for 24 and 57 days in 2024, and 2025 respectively. Other areas along the south coast, including Weymouth, West Bay, Torbay, and Dawlish, are likewise showing an increasing trend in elevated summer SSTs (Figure 1B). The observed increases in SST are consistent with reports from local fishermen noted above over the past decade and provide further evidence of the impacts of climate change in local coastal areas.

### *Vibrio* species establishing in the Teign Estuary

Over the past three decades, climate-driven increases in sea-surface temperature have been associated with the expansion of *Vibrio* species and a corresponding rise in vibriosis cases in both human populations and aquaculture systems [1, 91]. In the UK, ad hoc monitoring of waterways and shellfisheries conducted over the past 20 years has demonstrated increasing abundance and distribution of *Vibrio* species [13, 92–95]. Climate change has made coastal waters across many areas of southwest England increasingly suitable for the growth and establishment of both human and aquaculture-associated *Vibrio* species, potentially increasing vulnerability to the types of infections and disease outbreaks already observed internationally [13]. In the Teign Estuary at the Shaldon site, we have found a range of *Vibrio* species present at low abundances, including *V. parahaemolyticus, V. cholerae (*non-O1/non-O139*), V. alginolyticus, V. diabolicus*, *V. jasicida, V. aestuarianus, V. anguillarum, V. neptunius* and *V. pacinii,* which describes a strong and well-established *Vibrio* community presence in the waters. Furthermore, for *V. parahaemolyticus, V. alginolyticus* and *V. diabolicus* we found multiple distinct lineages of each *Vibrio* species, reflecting the diversity of these species present in the Teign Estuary. Despite the enormous global diversity of the *Vibrio* species detected here, four clonal groups were found in both 2024 and 2025 suggesting individual lineages of these species can persist in the Teign Estuary from year to year.

Although information on community cases of vibriosis remains limited, two *V. alginolyticus* infections were reported in Devon during the summer of 2025. These included a wound infection and a gastrointestinal infection in a patient with known exposure to freshwater and sewage, highlighting the potential for locally acquired infections (*Royal Devon & Exeter Foundation NHS Trust and Torbay and South Devon NHS Foundation Trust, unpublished data)*. *Vibrio* isolates from patients were not kept and thus we have no epidemiological linkage to our circulating environmental *V. alginolyticus* strains. In the UK, diagnostic testing only occurs routinely for *V. cholerae* and is usually undertaken only in patients presenting with gastrointestinal symptoms who have recently travelled to countries where cholera is endemic [96], this potentially limits the detection of locally acquired infections of *V. cholerae* and non-V. *cholerae* infections as environmental reservoirs become increasingly established in coastal waters. Consequentially, *Vibrio* infections are considered substantially underreported, underscoring the need for enhanced surveillance and a clearer understanding of the pathogen’s potential emergence in non-endemic regions such as the United Kingdom. Recently, reports of *Vibrio* infections in Ireland from patients lacking recent travel history have been reported [97]. Here authors, describe cases of otitis externa and skin/soft tissue infections in the summers of 2021(n=1) and 2022 (n=6) following exposure to seawater. Infections were linked to *V. alginolyticus* (n=4)*, V. diabolicus* (n=1)*, V. metschnikovii* (n=1) and non-O1/non-O139 *V. cholerae* (n=1). This recent published research along with the data we present in this study, is particularly relevant in the context of rapid climate warming in north-west Europe, which are influencing the distribution and abundance of marine *Vibrio* species.

Collectively, these observations suggest a potential expansion in both the seasonal duration and geographic range of suitable environmental conditions for *Vibrio* species. Accordingly, future research should prioritise monitoring *Vibrio* populations along the south coast of England, with particular emphasis on shellfish harvesting areas, in order to better characterise environmental drivers and associated public health risks.

### Environmental drivers of pathogenic *Vibrio* species

A storm overflow is a general term for an outlet that releases excess wastewater during heavy rainfall to prevent flooding of the sewer network. In 2024 and 2025, there were 1,437 and 1,446 storm overflow events, respectively, in the Teignmouth area of the UK. The majority of these events occurred during the winter months; however, between May and October, there were 538 events in 2024 and 48 events in 2025 in Teignmouth. The locations of the storm overflow releases are shown in Figure 1A. Enumeration of coliform bacteria, *Escherichia coli* and intestinal Enterococci (IE) can be used to indicate the presence of faecal contamination. The Environment Agency generally monitors bathing waters weekly during the bathing season. In our study, levels of *E. coli* and IE were measured monthly, while the Friends of the River Teign (FORT) also monitored water quality during the non-bathing season, generally every two weeks. Together, these monitoring programmes provide only a snapshot of water quality conditions and may not capture short-term pollution events.

A pollution incident is a broader term describing an event that causes environmental harm or breaches environmental regulations. This can include untreated sewage spills or storm overflow discharges that result in unacceptable environmental impacts. The length of time required for bathing waters to recover following a storm overflow event remains uncertain and may range from several hours to more than a day, depending on environmental conditions. Faster clearance of contamination is generally associated with high water exchange, such as strong tidal currents or wave action, which help disperse contaminated water. Biological filtration by shellfish may also contribute to improving water quality locally. However, where storm overflow discharges continue, spills are larger, or prolonged rainfall generates additional polluted runoff, contamination may persist for several days before water quality returns to safer levels. Despite considerable variability in composition, sewage effluent and surface runoff are often enriched in dissolved organic carbon (DOC; ∼40–80 mg/L), nitrogen (20–70 mg/L), and phosphate (4–8 mg/L), and may also contain heavy metals and residual concentrations of antibiotics [98–101]. Thus, sewage spills and storm overflows provide a nutrient influx that can impact aquatic ecosystems by stimulating the growth of native bacterial communities, including human-associated pathogens responsible for severe seafood-borne illness, such as *Vibrio* species.

We found in our study that untreated sewage proliferates the growth rate of different *Vibrio* species isolated from the Teign Estuary and that both sewage addition and temperature changes promote the growth of *V. cholerae* EXE 45/2024-E, *V. parahaemolyticus* EXE-19/2024-C, and *V. diabolicus* EXE 07/2023-A4 significantly. Previous studies have shown that sewage addition can increase the growth of *V. vulnificus* and promote biofilm formation by altering gene expression [102]. Studies have shown that the addition of sewage (50-100% treated sewage effluent) significantly increased the growth rates in *V. vulnificus* [102], while research by Conrad *et al.* found that as little as 1% raw sewage addition to estuarine water increased the *V. vulnificus* populations by 2-3 orders of magnitude [99]. This may pose an important risk to public health, particularly in coastal environments.

A review by Huijbers *et al.* in 2015 showed antimicrobial resistant (AMR) bacteria were detected in all publications investigating wastewater and included evidence for AMR transmission from wastewater to the environment [103–108]. Exposure to relevant sites for AMR and AMR genes (ARGs) includes beaches, recreational waters and shellfish, all of which have been documented as reservoirs of AMR-producing bacteria [104–109]. In this study, our preliminary scan of AMR bacteria and ARGs in the *Vibrio* community of water, sediment and various shellfish residing in the Teign Estuary showed the *Vibrio* strains to be multidrug resistant and to harbour an array of ARGs. Several AMRFinderPlus hits identified in this study likely represent components of the intrinsic resistome of *Vibrio* species rather than recently acquired resistance determinants. In particular, *tet(35)* was widely distributed among isolates from multiple species and has previously been proposed to constitute an intrinsic tetracycline resistance determinant in marine vibrios. Likewise, the *almEFG* operon detected in *V. cholerae* forms part of the lipid A modification pathway responsible for intrinsic polymyxin resistance in El Tor strains[110]. In contrast, genes detected only sporadically within the dataset, including *qnrC*, *fos* and *blaVHW*, may reflect acquired resistance events and warrant further investigation. The origin of AMR and ARG’s observed in these *Vibrio* species is unknown. The long-term historical mining that has taken place around the River Teign has been known to contribute to significant sources of zinc (Zn), cadmium (Cd), lead (Pb), Copper (Cu) and Barium (Ba) into the river [111, 112] and environmentally relevant levels of heavy metals can select for antimicrobial resistance [113, 114]. It is currently unknown if urban runoff, industrialised waste runoff, storm overflow discharges and other pollution incidents inputs into the Teign Estuary contribute to the acquisition and dissemination of AMR genes within the Teign Estuary *Vibrio* community.

Future studies should include sampling during and immediately following storm overflow events and/or periods of heavy rainfall. Such investigations would determine whether *Vibrio* abundances increase in response to these events, whether particular *Vibrio* species benefit from the sudden nutrient enrichment, how long any elevated increase in *Vibrio* levels persist, and whether the prevalence of AMR profiles and ARGs change following these pollution inputs. Addressing this knowledge gap will be essential for improving guidance on the safe management of bathing waters, including decisions on beach closures and reopening, during the summer period and for reducing potential public health risks.

## Methods and Materials

### Sample Collection and sea surface temperature monitoring

One sampling site was selected in the Teign Estuary at Shaldon Bridge and samples were collected monthly. Water temperature and pH was taken at the time of sampling using a pH and temperature probe (Hannah Instruments Ltd). Between August 2024 and September 2025, a HOBO MX2202 logger (Tempcon Instrumentation Ltd) was deployed in the Teign Estuary by attaching it to the rising chain of a mooring buoy. The loggers were approximately at 50cm depth at most states of the tide (except during the very lowest tides, where they would remain below the surface but at shallower depths. The loggers recorded the water temperature every 30 minutes, and from this, daily averages were calculated 9Figure 1C). A further four coastal locations were chosen for their proximity to Teignmouth, and SST data were taken from Directional Waverider buoys (DWR) MkIII (which record wave data every 30 minutes). Daily average temperatures were calculated, and the number of days where the daily average exceeded 18°C has been recorded in Figure 1B.

Every month, a grab sample consisting of 1L of seawater was collected. A sediment sample was collected from the top layer using a sterile 1L bottle attached to the end of a 3-metre sampling pole. The bottle was dragged with a backwards-forwards motion to collect around 400 mL of sediment, the sediment sample was left to settle, and any additional water was poured away, leaving the sediment only. Wild/non-depurated shellfish were collected each month when available and consisted of Pacific oysters (*Magallana gigas*), blue mussels (*Mytilus edulis*), cockles (*Cerastoderma edule*) and surf clams (*Spisula solidissima*). Cockles were predominantly available all year round. All samples were held between 4 and 8°C during transit and, on arrival in the laboratory, were placed in the fridge until analysis.

### Microbiological Analysis of Water, Sediment, and Shellfish Samples

All samples were analysed within 24 h of collection and according to ISO 21872-1 with minor modifications. For water samples, 100 µl was spread directly onto Membrane Lactose Glucuronide Agar (MLGA) for the differentiation of *Escherichia coli* and other coliforms in water, and on thiosulphate citrate bile sucrose agar (TCBS) plates for the differentiation of *Vibrio* species.

Additionally, 25ml of water sample was added to 225 ml of alkaline salt peptone water (ASPW). For sediment samples, 25 g of sediment was added to 225ml ASPW and shaken by hand, after which 100 µl was spread directly onto MLGA and TCBS plates as above. A total of 25 g of shellfish meat (taken from a minimum of 8-10 oysters, 12-15 mussels, 8-10 clams or 15 cockles) was disrupted using a stomacher and added to 225 ml of ASPW, after which 100 µl was spread directly onto MLGA and TCBS plates. Sequential log dilutions of samples were carried out in APSW to a maximum of 10^-2^, and 100 µl was spread onto the surface of MLGA and TCBS plates to determine direct enumeration of *E. coli,* other coliforms and *Vibrio* species. All enrichment broths were then incubated at 41 °C for 6 h for an enrichment step, after which a 5 µl loopful was taken from directly below the surface of the broth and streaked onto TCBS plates. Up to 5 colonies, including both sucrose-negative (green) and sucrose-positive (yellow) morphotypes, were selected and subcultured onto marine agar (SLS labs) and *Vibrio* chromID (VID) agar (BioMérieux) where fewer than five colonies were present, all colonies were included. All MLGA plates were incubated at 30°C for 4 h before incubation at 37°C for 18-24 h. All TCBS and VID plates were incubated at 37 °C for 18-24 h. All Marine agar was incubated at 30 °C for 18-24 h.

Presumptive colonies were identified as *V. parahaemolyticus* if they met the following criteria: non-sucrose (green) on TCBS plates and pink on VID agar and positive for the enzyme cytochrome c oxidase (oxidase test). Further identification by API 20E strips (BioMérieux) was also carried out on both sucrose negative and sucrose positive colonies. All biochemically identified *V. parahaemolyticus* strains were further analysed by PCR amplification using the species target toxR [30]. All toxR-positive strains were tested for PCR amplification of *tdh* and *trh* genes using primers designed by Tada *et al* [31]. All sucrose-negative colonies that were also negative for *V. parahaemolyticus* PCR tests were checked for *V. vulnificus* identification using PCR amplification using the species target *vvh* [115].

### Additional testing by Friends of the River Teign (FORT)

The local Friends of the River Teign (FORT) group collected water samples on randomised days every week at the Ferry Shelter Shaldon Beach in the Teign Estuary. The samples were taken at a minimum 30cm depth in a minimum of 1m depth of water, using a sterilised container positioned more than a metre away from the sample collector. Samples are taken on a transit line between two fixed marks on Shaldon Beach. This ensured that the samples were taken at a fixed location relative to any pollution sources. However, the exact position along the transit line was determined by the state of the tide when the sample was taken. At low water, the location is approximately at clouding.secretly.front (what3words). After collection, the samples were sent to Eurofins for commercial testing.

### DNA sequencing

Fifty-Three strains were selected for whole genome sequencing. Bacterial cells were harvested, and genomic DNA was extracted using the Wizard Genomic DNA Purification Kit (Promega). Library preparation was carried out by Novogene Europe and Exeter DNA Sequencing Service. This study used bacterial genome sequences assembled from Illumina (short) and / or ONT (long) sequencing reads, deposited in the Sequence Read Archive (SRA) [116] under BioProject PRJNA1453982. The genome sequencing was performed in several batches between May 2024 and December 2025. The first batch was sequenced using the Illumina NovaSeq 6000 by Novogene. Subsequent batches were sequenced by the University of Exeter Sequencing Facility using the ONT PromethION only, or a combination of PromethION plus Illumina NovaSeq X. Supplementary Table S4 lists the sequencing method(s) applied to each isolate.

### Quality-based filtering of sequence reads

Short (Illumina) reads were trimmed using TrimGalore [117]0.6.10 with command-line options “-q 30” and “--paired”. Long, ONT reads were filtered using Filtlong 0.2.1 [118]. The command line for each ONT FASTQ file was: “filtlong --min_length 1000 --keep_percent 95 fastq.gz | gzip > filtlong.fastq.gz”.

### *De-novo* genome assembly

We used assembly tools appropriate to the type of sequence data to be assembled. Specifically, for assembling genomes from only short reads, we used Unicycler 0.5.1 [119]. For hybrid assembly of trimmed short reads combined with filtered long reads, we used Unicycler 0.5.1[119]. The command line for each genome assembly was “unicycler -1 trimmed-reads-1.fq.gz -2 trimmed-reads-2.fq.gz -l filtlong.fastq.gz -o out.unicycler”. For assembling genomes from long reads only, we used Hybracter v 0.11.2 [120] with command line “ hybracter long -i hybracter.input.csv”. The Hybracter.input.csv file is available from GitHub at https://github.com/davidjstudholme/vibrio/raw/refs/heads/main/12_hybracter/P0213_250813 _11813/02_hybracter/hybracter.input.csv. We used non-filtered reads, because the Hybracter pipeline contains its own quality-control steps [120]. Assembly statistics were assessed using Quast 5.0.2 [121] and CheckM v1.2.4 [122] and isolates failing QC were removed from the study. These same metrics were also automatically calculated by the NCBI after we submitted the assemblies to GenBank [123] via the NCBI’s Submission Portal [124]. The assembled genome sequences were annotated after submission by the NCBI using PGAP [125].

### Species identification

Reads were classified to species level using Sylph v 2.3 [126]. To infer taxonomic species from assembled genome sequences, we used the Type Strain Genome Server (TYGS) [56]. To calculate average nucleotide identity (ANI) between genomes of our isolates versus type strains, we used fastANI 1.33 [127].

### Multi-locus sequence typing

Multi-locus sequence typing was performed on each isolate using MLST 2.33.1 [128] [129] with automatic typing scheme selection. The MLST software made use of the PubMLST website (https://pubmlst.org/) [128]. Allele combinations detected with no associated ST name in the typing schemes were given a novel sequence type number (NSTXX).

### Phylogenomic overview of *Vibrio* genus

We placed the newly sequenced genomes into the phylogenetic context of type strains of *Vibrio* species. To achieve this, we generated approximate maximum likelihood trees based on core genome sequences, using PhaME 1.0.3 [130] and FastTree 2 [131]. The command line for each PhaME tree was “phame config-file.ctl”. The configuration files can be found at https://github.com/davidjstudholme/vibrio/tree/main/01_PhaME/05_phame.

### Phylogenomic analysis of Vibrio parahaemolyticus and V. diabolicus

Publicly accessible genome assemblies of *V. parahaemolyticus*, *V. diabolicus*, *V. alginolyticus* and *V. antiquarius* were downloaded from the NCBI using NCBI datasets v18.28.0 [132]. For *V. diabolicus*, *V. alginolyticus* and *V. antiquarius* genomes were combined with study isolates and a phylogenetic tree was constructed using MashTree v 1.4.6 [133] with parameter –mindepth 0. For *V. parahaemolyticus*, before building a phylogeny in a similar manner, 14,702 publicly accessible isolates that were not isolated in the UK were dereplicated by calculating pairwise MinHash distances with Mash v 2.3 [134] and retaining one representative genome per 2129 clusters within 0.5 MinHash distance. Eight outlier genomes were excluded based on manual inspection of preliminary trees. Phylogenies were visualised using Microreact v 293 [135] and R version 4.4.3 [136].

### Prediction of virulence factors and anti-microbial resistance (AMR) markers

To identify virulence genes, we scanned genome assemblies against VFDB [137], using ABRicate 1.4.0 [138].AMR genes were detected in assembled genome sequences using AMRFinderPlus 4.2.7 [87].

### PCR detection of *V. diabolicus* strains

Supplementary Table S5 shows the primers and conditions designed to detect hypothetical protein (ACY49842.1), hydroxyectoine utilisation dehydratase (ACY53324.1), ornithine cyclodeaminase (ACY53325.1) and phosphoesterase (ACY53763.1) using PCR to determine if they could be used to detect *V. diabolicus* strains.

### Infection of Galleria mellonella larvae

*Galleria mellonella* larvae were purchased from UK Wax Worms Ltd. Larvae weighing between 0.2 and 0.35 g were chosen for experiments. For each experiment, a total of ten larvae were used per strain to be tested. The larvae were infected by micro-injection (Hamilton Ltd) into the right foremost proleg with 10^2^-10^3^ CFU per larva of each strain to be tested in 10 µl volume, which had been grown in Marine Broth at 30 °C and washed twice in PBS. Bacterial cell counts were carried out by plating serial dilutions of the inoculum onto Marine agar. For control purposes, ten larvae were inoculated with PBS. The larvae were incubated at 37°C, and survival was recorded for all strains after 24 h and 48 h. Larvae were scored as dead when they ceased moving or failed to respond when gently manipulated with a pipette tip. Observation findings were also recorded if larvae changed colour from their normal pale cream colouration to brown or black, indicative of melanisation.

### Co-culture with sewage

Primary untreated (crude) influent sewage was collected from a local Wastewater Treatment Plant (WWTP) in August 2025. Sewage was transported back to the laboratory in a cool box and then filter sterilised through a 0.2μm filter. To ensure sterility, 100μl aliquots of filtered sewage were spread onto Luria-Bertani (LB) agar plates and incubated at 37°C for 24h. If no colonies were recorded after 24h, sewage was recorded as sterile. Aliquots of filtered sewage (50ml falcon tubes) were stored in a -20°C freezer and thawed immediately prior to use. Filtered sewage was then co-cultured with bacteria in MSYE (modified yeast seawater extract).

Four *Vibrio* strains isolated from the Teign Estuary were selected to determine the impacts of wastewater influent on their growth: *V. parahaemolyticus* EXE-19/2024-C, *V. alginolyticus* EXE-09/2024-B, *V. diabolicus* EXE07/2023-A4, and *V. cholerae* EXE-45/2025-E were selected. Overnight cultures of selected strains were grown in LB broth at 37°C. Starting cultures of each strain were achieved by diluting to 0.02 OD590nm in Modified Seawater Yeast Extract (MSYE) [139]. A dilution series of MYSE:sewage across a 96-well plate was set up to give 50, 25, 12.5, and 0 % sewage concentrations. Aliquots of 100μl of starting cultures of 0.02 OD strains in MYSE were then added to wells, giving a final OD of 0.01. Control wells contained either sewage only, MSYE only, or MSYE and bacteria (0% sewage). The optical density at 590nm (OD590) of each replicate was measured every 30 minutes for 12 hours using a Tecan Infinite®200PRO. Experiments were carried out at 25 and 30°C to reflect either environmentally relevant temperatures or optimum growth temperatures for *Vibrio* strains [140]. Each plate consisted of three replicates per strain (technical repeats), and each experiment was repeated three times for each temperature (experimental repeats). Raw data files were analysed in Excel, RStudio and GraphPad Prism. Background OD was calculated from MSYE only wells and used to calculate the final OD595 from experimental wells. Growth data and growth rate (r) were analysed and calculated in RStudio (R version 4.5.0) using the growthcurver package [141]. Further statistical analysis was performed in GraphPad Prism.

### Working out the Minimum inhibitory concentration of Various *Vibrio* strains

The minimum concentration of antibiotic required to inhibit the growth of 13 *Vibrio* strains was tested using broth microdilution according to established protocols [36]. The antibiotics tested were ampicillin, kanamycin, chloramphenicol, trimethoprim, ceftazidime, ciprofloxacin, doxycycline, and gentamicin. Stock antibiotics were aliquoted and stored at –20°C in appropriate solvents. Overnight cultures of *Vibrio* strains were grown in LB broth at 37°C (except *V. aestuarianus* EXE 13/2025-H at 30°C). Two-fold serial dilutions of antibiotics ranged from 1024 μg/mL to 1 μg/mL in 96-well plates, inoculated to a final bacterial density of ∼10^6^ CFU ml^-1^ (OD_595nm = 0.01). Each plate contained triplicate technical repeats (except *V. parahaemolyticus* EXE 29/2025-HH, which was done in duplicate), no-antibiotic growth controls, and no-bacteria sterility controls. Plates were incubated static at respective temperatures for 22–24 hours. Growth was quantified using a Tecan microplate reader measuring OD 595nm. MIC was defined as the lowest antibiotic concentration inhibiting visible growth. The resistance or susceptibility is determined by using the MIC clinical breakpoints, which were found in previous research [142, 143]. The multiple antibiotic resistance (MAR) index was calculated for each strain by dividing the number of antibiotics they were resistant to by the overall number of antibiotics tested.

## Acknowledgments

Many thanks go to members of the Friends of the River Teign (FORT) group including Stuart Reynolds, Alec Collyer, Maurizio Pupi, Rob Parsons and Betina Winkler for their knowledge regarding the River Teign, helping with collecting logger data and aiding with sampling and general enthusiasm for our research. Thank you to Megan Dymond for assisting us in sampling during the summer of 2025. Thank you to Marina Morgan and Debbie Beer at Royal Devon and Exeter Hospital and Giuseppe Pichierri, Alexandria Willis, and Cheryl Bailiss at Torbay Hospital. Thank you to South West Water for proving us with primary untreated (crude) influent sewage. This project was generously supported by NE/X018032/1 from the Natural Environment Research Council. NC’s work was supported by BBSRC grant BB/Z515139/1. We acknowledge the University of Exeter Sequencing Service for their assistance with whole genome sequencing. This project utilised equipment funded by the UK Medical Research Council (MRC) Clinical Research Infrastructure Initiative (award number MR/M008924/1). This project utilised equipment funded by the BBSRC ALERT mid-range equipment 2024 award (BB/Z515942/1).

For the purpose of open access, the author has applied a Creative Commons Attribution (CC BY) licence to any Author Accepted Manuscript version arising from this submission.

## Author Contributions

HB and SW conceived and designed the experiments. HB, AF, IA, MB, SR and SW performed the experiments. NC and DJS carried out genome assembly and comparative genomics. HB, NC, AF, MB, DJS and SW analysed the data. SW wrote the first draft of the paper and HB, NC, AF, IF, IA, MB, DJS and SW contributed to subsequent versions. All Authors read and approved the final manuscript.

## Supplementary data

Supplementary Table S1: Water, Sediment, Shellfish and Seaweed samples tested in this study between 2023 and 2025.

Supplementary Table S2: Dose range injected to *G. mellonella*

Supplementary Table S3: Antibiotic susceptibilities.

Supplementary Table S4. Genome sequencing for each of the *Vibrio* isolates sequenced in this study.

Supplementary Table S5: PCR primers and conditions for identifying V. diabolicus isolates.

Supplementary Figure S2. Mash tree of 2229 *Vibrio parahaemolyticus* isolates including 12 study isolates.

Supplementary Figure S3. Mash tree of isolates submitted to the NCBI as *V. alginolyticus* (n=488), *V. diabolicus* (n=158) and *V. antiquarius* (n=19) isolates including 22 study isolates.

Supplementary Figure S3: Source of growth rate variation

## References

1. Trinanes J, Martinez-Urtaza J. Future scenarios of risk of Vibrio infections in a warming planet: a global mapping study. Lancet Planet Health. 2021;5(7):e426–e35; doi: 10.1016/s2542-5196(21)00169-8.

2. Agency UHS: Gastrointestinal infections in England: 2022 to 2024. In.; March 2026.

3. Baker-Austin C, Jenkins C, Dadzie J, Mestanza O, Delgado E, Powell A, et al. Genomic epidemiology of domestic and travel-associated Vibrio parahaemolyticus infections in the UK, 2008–2018. Food Control. 2020;115:107244; doi: 10.1016/j.foodcont.2020.107244.

4. Poh CJ, Greig DR, Rodwell EV, Jenkins C. Public health surveillance of Vibrio cholerae in travellers returning to the United Kingdom. J Med Microbiol. 2026;75(2); doi: 10.1099/jmm.0.002121.

5. DiSalvo LH, Blecka J, Zebal R. Vibrio anguillarum and larval mortality in a California coastal shellfish hatchery. Appl Environ Microbiol. 1978;35(1):219–21; doi: 10.1128/aem.35.1.219-221.1978.

6. Lacoste A, Jalabert F, Malham S, Cueff A, Gélébart F, Cordevant C, et al. A Vibrio splendidus strain is associated with summer mortality of juvenile oysters Crassostrea gigas in the Bay of Morlaix (North Brittany, France). Dis Aquat Organ. 2001;46(2):139–45; doi: 10.3354/dao046139.

7. Garnier M, Labreuche Y, Garcia C, Robert M, Nicolas JL. Evidence for the involvement of pathogenic bacteria in summer mortalities of the Pacific oyster Crassostrea gigas. Microb Ecol. 2007;53(2):187–96; doi: 10.1007/s00248-006-9061-9.

8. Malham SK, Cotter E, O’Keeffe S, Lynch S, Culloty SC, King JW, et al. Summer mortality of the Pacific oyster, Crassostrea gigas, in the Irish Sea: The influence of temperature and nutrients on health and survival. Aquaculture. 2009;287(1):128–38; doi: 10.1016/j.aquaculture.2008.10.006.

9. Le Roux F, Gay M, Lambert C, Waechter M, Poubalanne S, Chollet B, et al. Comparative analysis of Vibrio splendidus-related strains isolated during Crassostrea gigas mortality events. Aquat Living Resour. 2002;15(4):251–8; doi: 10.1016/S0990-7440(02)01176-2.

10. de Lorgeril J, Escoubas JM, Loubiere V, Pernet F, Le Gall P, Vergnes A, et al. Inefficient immune response is associated with microbial permissiveness in juvenile oysters affected by mass mortalities on field. Fish Shellfish Immunol. 2018;77:156–63; doi: 10.1016/j.fsi.2018.03.027.

11. Motes ML, DePaola A, Cook DW, Veazey JE, Hunsucker JC, Garthright WE, et al. Influence of water temperature and salinity on Vibrio vulnificus in Northern Gulf and Atlantic Coast oysters (Crassostrea virginica). Appl Environ Microbiol. 1998;64(4):1459–65; doi: 10.1128/AEM.64.4.1459-1465.1998.

12. Yoon KS, Min KJ, Jung YJ, Kwon KY, Lee JK, Oh SW. A model of the effect of temperature on the growth of pathogenic and nonpathogenic Vibrio parahaemolyticus isolated from oysters in Korea. Food Microbiol. 2008;25(5):635–41; doi: 10.1016/j.fm.2008.04.007.

13. Harrison J, Nelson K, Morcrette H, Morcrette C, Preston J, Helmer L, et al. The increased prevalence of Vibrio species and the first reporting of Vibrio jasicida and Vibrio rotiferianus at UK shellfish sites. Water Res. 2022;211:117942; doi: 10.1016/j.watres.2021.117942.

14. 2009 VSDCICL. The River Teign. doi: https://www.visitsouthdevon.co.uk/things-to-do/the-river-teign-p1541153.

15. Coles T. Impacts of climate change on tourism and marine recreation MCCIP Science 2020:593–615; doi: 10.14465/2020.arc25.tre

16. Society RM. UK seasonal weather summary Summer 2024. Weather. 2024;79(10):325-; doi: 10.1002/wea.7625.

17. Randa MA, Polz MF, Lim E. Effects of temperature and salinity on Vibrio vulnificus population dynamics as assessed by quantitative PCR. Appl Environ Microbiol. 2004;70(9):5469–76; doi: 10.1128/AEM.70.9.5469-5476.2004.

18. England. NNoRCMPo. https://coastalmonitoring.org/.

19. Affairs DfEFR. Bathing Water Quality: Swimfo: Find a bathing water. 2026.

20. Department for Environment FRA. ENV17 - Bathing water quality: additional datasets. 2025.

21. Water SW. Escherichia Coli and Intestinal Enterococci. 2026.

22. Lees D. Viruses and bivalve shellfish. Int J Food Microbiol. 2000;59(1-2):81–116; doi: 10.1016/s0168-1605(00)00248-8.

23. Oliver JD, Warner RA, Cleland DR. Distribution of Vibrio vulnificus and other lactose-fermenting vibrios in the marine environment. Appl Environ Microbiol. 1983;45(3):985–98; doi: 10.1128/aem.45.3.985-998.1983.

24. Froelich B, Oliver JD. The interactions of Vibrio vulnificus and the oyster Crassostrea virginica. Microb Ecol. 2013;65(4):807–16; doi: 10.1007/s00248-012-0162-3.

25. Malham SK, Taft H, Farkas K, Ladd CJT, Seymour M, Robins PE, et al. Multi-scale influences on Escherichia coli concentrations in shellfish: From catchment to estuary. Environmental Pollution. 2025;366:125476; doi: 10.1016/j.envpol.2024.125476.

26. Malham SK, Rajko-Nenow P, Howlett E, Tuson KE, Perkins TL, Pallett DW, et al. The interaction of human microbial pathogens, particulate material and nutrients in estuarine environments and their impacts on recreational and shellfish waters. Environ Sci Process Impacts. 2014;16(9):2145–55; doi: 10.1039/c4em00031e.

27. Kobayashi T, Enomoto S, Sakazaki R, Kuwahara S. [a New Selective Isolation Medium for the Vibrio Group; on a Modified Nakanishi’s Medium (Tcbs Agar Medium)]. Nihon Saikingaku Zasshi. 1963;18:387–92; doi: 10.3412/jsb.18.387.

28. Reilly GD, Reilly CA, Smith EG, Baker-Austin C. Vibrio alginolyticus-associated wound infection acquired in British waters, Guernsey, July 2011. Euro Surveill. 2011;16(42).

29. JB OJaK. Vibrio Species,” In: M. P. Doyle, L. R. Beuchat and T. J. Montville, Eds., Food Microbiology: Fundamentals and Frontiers,. 1997:228-64.

30. Kim YB, Okuda J, Matsumoto C, Takahashi N, Hashimoto S, Nishibuchi M. Identification of Vibrio parahaemolyticus strains at the species level by PCR targeted to the toxR gene. J Clin Microbiol. 1999;37(4):1173–7.

31. Tada J, Ohashi T, Nishimura N, Shirasaki Y, Ozaki H, Fukushima S, et al. Detection of the thermostable direct hemolysin gene (tdh) and the thermostable direct hemolysin-related hemolysin gene (trh) of Vibrio parahaemolyticus by polymerase chain reaction. Mol Cell Probes. 1992;6(6):477–87.

32. Albert MJ, Siddique AK, Islam MS, Faruque AS, Ansaruzzaman M, Faruque SM, et al. Large outbreak of clinical cholera due to Vibrio cholerae non-O1 in Bangladesh. Lancet. 1993;341(8846):704; doi: 10.1016/0140-6736(93)90481-u.

33. Large epidemic of cholera-like disease in Bangladesh caused by Vibrio cholerae O139 synonym Bengal. Cholera Working Group, International Centre for Diarrhoeal Diseases Research, Bangladesh. Lancet. 1993;342(8868):387–90.

34. Huq A, Parveen S, Qadri F, Sack DA, Colwell RR. Comparison of Vibrio cholerae serotype 01 strains isolated from patients and the aquatic environment. The Journal of tropical medicine and hygiene. 1993;96(2):86–92.

35. Hall RH, Khambaty FM, Kothary MH, Keasler SP, Tall BD. Vibrio cholerae non-O1 serogroup associated with cholera gravis genetically and physiologically resembles O1 E1 Tor cholera strains. Infect Immun. 1994;62(9):3859–63; doi: 10.1128/iai.62.9.3859-3863.1994.

36. Herrington DA, Hall RH, Losonsky G, Mekalanos JJ, Taylor RK, Levine MM. Toxin, toxin-coregulated pili, and the toxR regulon are essential for Vibrio cholerae pathogenesis in humans. J Exp Med. 1988;168(4):1487–92; doi: 10.1084/jem.168.4.1487.

37. Waldor MK, Mekalanos JJ. Lysogenic conversion by a filamentous phage encoding cholera toxin. Science. 1996;272(5270):1910–4; doi: 10.1126/science.272.5270.1910.

38. Singh DV, Matte MH, Matte GR, Jiang S, Sabeena F, Shukla BN, et al. Molecular analysis of Vibrio cholerae O1, O139, non-O1, and non-O139 strains: clonal relationships between clinical and environmental isolates. Appl Environ Microbiol. 2001;67(2):910–21; doi: 10.1128/aem.67.2.910-921.2001.

39. T S. BRicate: Mass screening of contigs for antimicrobial and virulence genes. GitHub https://githubcom/tseemann/abricate. 2020; doi: https://github.com/tseemann/abricate.

40. Altschul SF, Gish W, Miller W, Myers EW, Lipman DJ. Basic local alignment search tool. J Mol Biol. 1990;215(3):403–10; doi: 10.1016/S0022-2836(05)80360-2.

41. Dutta D, Chowdhury G, Pazhani GP, Guin S, Dutta S, Ghosh S, et al. Vibrio cholerae non-O1, non-O139 serogroups and cholera-like diarrhea, Kolkata, India. Emerg Infect Dis. 2013;19(3):464–7; doi: 10.3201/eid1903.121156.

42. Chatterjee S, Ghosh K, Raychoudhuri A, Chowdhury G, Bhattacharya MK, Mukhopadhyay AK, et al. Incidence, virulence factors, and clonality among clinical strains of non-O1, non-O139 Vibrio cholerae isolates from hospitalized diarrheal patients in Kolkata, India. J Clin Microbiol. 2009;47(4):1087–95; doi: 10.1128/jcm.02026-08.

43. Rodríguez JY, Duarte C, Rodríguez GJ, Montaño LA, Benítez-Peñuela MA, Díaz P, et al. Bacteremia by non-O1/non-O139 Vibrio cholerae: Case description and literature review. Biomedica. 2023;43(3):323–9; doi: 10.7705/biomedica.6716.

44. Zmeter C, Tabaja H, Sharara AI, Kanj SS. Non-O1, non-O139 Vibrio cholerae septicemia at a tertiary care center in Beirut, Lebanon; a case report and review. J Infect Public Health. 2018;11(5):601–4; doi: 10.1016/j.jiph.2018.01.001.

45. Gherlan GS, Lazar DS, Florescu SA, Dirtu RM, Codreanu DR, Lupascu S, et al. Non-toxigenic Vibrio cholerae - just another cause of vibriosis or a potential new pandemic? Arch Clin Cases. 2025;12(1):5–16; doi: 10.22551/2025.46.1201.10305.

46. Turner JW, Tallman JJ, Macias A, Pinnell LJ, Elledge NC, Nasr Azadani D, et al. Comparative Genomic Analysis of Vibrio diabolicus and Six Taxonomic Synonyms: A First Look at the Distribution and Diversity of the Expanded Species. Front Microbiol. 2018;9:1893; doi: 10.3389/fmicb.2018.01893.

47. Ren C, Hu C, Jiang X, Sun H, Zhao Z, Chen C, et al. Distribution and pathogenic relationship of virulence associated genes among Vibrio alginolyticus from the mariculture systems. Mol Cell Probes. 2013;27(3-4):164–8; doi: 10.1016/j.mcp.2013.01.004.

48. Austin B, Stuckey LF, Robertson PAW, Effendi I, Griffith DRW. A probiotic strain of Vibrio alginolyticus effective in reducing diseases caused by Aeromonas salmonicida, Vibrio anguillarum and Vibrio ordalii. Journal of fish diseases. 1995;18(1):93–6; doi: 10.1111/j.1365-2761.1995.tb01271.x.

49. Vandenberghe J, Li Y, Verdonck L, Li J, Sorgeloos P, Xu HS, et al. Vibrios associated with Penaeus chinensis (Crustacea: Decapoda) larvae in Chinese shrimp hatcheries. Aquaculture. 1998;169(1):121–32; doi: 10.1016/S0044-8486(98)00319-6.

50. Skliros D, Kostakou M, Kokkari C, Tsertou MI, Pavloudi C, Zafeiropoulos H, et al. Unveiling Emerging Opportunistic Fish Pathogens in Aquaculture: A Comprehensive Seasonal Study of Microbial Composition in Mediterranean Fish Hatcheries. Microorganisms. 2024;12(11); doi: 10.3390/microorganisms12112281.

51. Gomez-Leon J, Villamil L, Lemos ML, Novoa B, Figueras A. Isolation of Vibrio alginolyticus and Vibrio splendidus from aquacultured carpet shell clam (Ruditapes decussatus) larvae associated with mass mortalities. Appl Environ Microbiol. 2005;71(1):98–104; doi: 10.1128/AEM.71.1.98-104.2005.

52. Raguenes G, Christen R, Guezennec J, Pignet P, Barbier G. Vibrio diabolicus sp. nov., a new polysaccharide-secreting organism isolated from a deep-sea hydrothermal vent polychaete annelid, Alvinella pompejana. Int J Syst Bacteriol. 1997;47(4):989–95; doi: 10.1099/00207713-47-4-989.

53. Sayers EW, Cavanaugh M, Clark K, Ostell J, Pruitt KD, Karsch-Mizrachi I. GenBank. Nucleic Acids Res. 2020;48(D1):D84–D6; doi: 10.1093/nar/gkz956.

54. Hasan NA, Grim CJ, Lipp EK, Rivera IN, Chun J, Haley BJ, et al. Deep-sea hydrothermal vent bacteria related to human pathogenic Vibrio species. Proc Natl Acad Sci U S A. 2015;112(21):E2813–9; doi: 10.1073/pnas.1503928112.

55. NCBI. https://www.ncbi.nlm.nih.gov/datasets/genome/?taxon=150340.

56. Freese HM, Meier-Kolthoff JP, Sardà Carbasse J, Afolayan Ayorinde O, Göker M. TYGS and LPSN in 2025: a Global Core Biodata Resource for genome-based classification and nomenclature of prokaryotes within DSMZ Digital Diversity. Nucleic Acids Res. 2026;54(D1):D884–D91; doi: 10.1093/nar/gkaf1110.

5 7. NCBI. https://www.ncbi.nlm.nih.gov/datasets/genome/?taxon=2527672.

58. Ghosh A, Bhadury P. Vibrio chemaguriensis sp. nov., from Sundarbans, Bay of Bengal. Curr Microbiol. 2019;76(10):1118–27; doi: 10.1007/s00284-019-01731-7.

59. Yoshizawa S, Tsuruya Y, Fukui Y, Sawabe T, Yokota A, Kogure K, et al. Vibrio jasicida sp. nov., a member of the Harveyi clade, isolated from marine animals (packhorse lobster, abalone and Atlantic salmon). Int J Syst Evol Microbiol. 2012;62(Pt 8):1864–70; doi: 10.1099/ijs.0.025916-0.

60. Diggles BK, Moss GA, Carson J, Anderson CD. Luminous vibriosis in rock lobster Jasus verreauxi (Decapoda: Palinuridae) phyllosoma larvae associated with infection by Vibrio harveyi. Dis Aquat Organ. 2000;43(2):127–37; doi: 10.3354/dao043127.

61. Balebona MC, Krovacek K, Moriñigo MA, Mansson I, Faris A, Borrego JJ. Neurotoxic effect on two fish species and a PC12 cell line of the supernate of Vibrio alginolyticus and Vibrio anguillarum. Veterinary microbiology. 1998;63(1):61–9; doi: 10.1016/S0378-1135(98)00227-2.

62. Naka H, Crosa JH. Genetic Determinants of Virulence in the Marine Fish Pathogen Vibrio anguillarum. Fish Pathol. 2011;46:1–10; doi: 10.3147/jsfp.46.1.

63. Crosa JH, Hodges LL, Schiewe MH. Curing of a plasmid is correlated with an attenuation of virulence in the marine fish pathogen Vibrio anguillarum. Infect Immun. 1980;27(3):897–902; doi: 10.1128/iai.27.3.897-902.1980.

64. Auguste M, Leonessi M, Oliveri C, Balbi T, Rahman FU, Vezzulli L, et al. Deciphering host-pathogen dynamics in Mytilus galloprovincialis: immune responses to multiple infections with bacterial isolates from bivalve mortality outbreaks. Fish Shellfish Immunol. 2026;169:111064; doi: 10.1016/j.fsi.2025.111064.

65. Lupo C, Dutta BL, Petton S, Ezanno P, Tourbiez D, Travers MA, et al. Spatial epidemiological modelling of infection by Vibrio aestuarianus shows that connectivity and temperature control oyster mortality. Aquacult Env Interac. 2020;12:511–27; doi: 10.3354/aei00379.

66. Parizadeh L, Tourbiez D, Garcia C, Haffner P, Dégremont L, Le Roux F, et al. Ecologically realistic model of infection for exploring the host damage caused by Vibrio aestuarianus. Environ Microbiol. 2018;20(12):4343–55; doi: 10.1111/1462-2920.14350.

67. Coyle NM, O’Toole C, Thomas JCL, Ryder D, Feil EJ, Geary M, et al. Vibrio aestuarianus clade A and clade B isolates are associated with Pacific oyster (Magallana gigas) disease outbreaks across Ireland. Microb Genom. 2023;9(8); doi: 10.1099/mgen.0.001078.

68. Goudenège D, Travers MA, Lemire A, Petton B, Haffner P, Labreuche Y, et al. A single regulatory gene is sufficient to alter Vibrio aestuarianus pathogenicity in oysters. Environ Microbiol. 2015;17(11):4189–99; doi: 10.1111/1462-2920.12699.

69. Garnier M, Labreuche Y, Nicolas JL. Molecular and phenotypic characterization of Vibrio aestuarianus subsp. francensis subsp. nov., a pathogen of the oyster Crassostrea gigas. Syst Appl Microbiol. 2008;31(5):358–65; doi: 10.1016/j.syapm.2008.06.003.

70. Prado S, Romalde JL, Montes J, Barja JL. Pathogenic bacteria isolated from disease outbreaks in shellfish hatcheries. First description of Vibrio neptunius as an oyster pathogen. Dis Aquat Organ. 2005;67(3):209–15; doi: 10.3354/dao067209.

71. Dubert J, Barja JL, Romalde JL. New Insights into Pathogenic Vibrios Affecting Bivalves in Hatcheries: Present and Future Prospects. Front Microbiol. 2017;8:762; doi: 10.3389/fmicb.2017.00762.

72. Galvis F, Ageitos L, Rodríguez J, Jiménez C, Barja JL, Lemos ML, et al. Vibrio neptunius Produces Piscibactin and Amphibactin and Both Siderophores Contribute Significantly to Virulence for Clams. Front Cell Infect Microbiol. 2021;11:750567; doi: 10.3389/fcimb.2021.750567.

73. Hu R-G, Yang L, Wang L-Y, Yang Y-L, Li H-J, Yang B-T, et al. Unveiling the pathogenic and multidrug-resistant profiles of Vibrio alfacsensis: A potential identified threat in turbot (Scophthalmus maximus) aquaculture. J Hazard Mater. 2024;479:135729; doi: 10.1016/j.jhazmat.2024.135729.

74. Gomez-Gil B, Thompson FL, Thompson CC, Swings J. Vibrio pacinii sp. nov., from cultured aquatic organisms. Int J Syst Evol Microbiol. 2003;53(Pt 5):1569–73; doi: 10.1099/ijs.0.02670-0.

75. Birkbeck TH, Treasurer JW. Vibrio splendidus, Vibrio ichthyoenteri and Vibrio pacinii isolated from the digestive tract microflora of larval ballan wrasse, Labrus bergylta Ascanius, and goldsinny wrasse, Ctenolabrus rupestris (L.). J Fish Dis. 2014;37(1):69–74; doi: 10.1111/jfd.12116.

76. Zanetti S, Deriu A, Volterra L, Falchi MP, Molicotti P, Fadda G, et al. Virulence factors in Vibrio alginolyticus strains isolated from aquatic environments. Ann Ig. 2000;12(6):487–91.

77. Balebona MC, Andreu MJ, Bordas MA, Zorrilla I, Moriñigo MA, Borrego JJ. Pathogenicity of *Vibrio alginolyticus*for Cultured Gilt-Head Sea Bream (*Sparus aurata*L.). Applied and environmental microbiology. 1998;64(11):4269–75; doi: doi:10.1128/AEM.64.11.4269-4275.1998.

78. Sanches-Fernandes GMM, Sá-Correia I, Costa R. Vibriosis Outbreaks in Aquaculture: Addressing Environmental and Public Health Concerns and Preventive Therapies Using Gilthead Seabream Farming as a Model System. Front Microbiol. 2022;13:904815; doi: 10.3389/fmicb.2022.904815.

79. Wagley S, Borne R, Harrison J, Baker-Austin C, Ottaviani D, Leoni F, et al. Galleria mellonella as an infection model to investigate virulence of Vibrio parahaemolyticus. Virulence. 2018;9(1):197–207; doi: 10.1080/21505594.2017.1384895.

80. Sathyamoorthy V, Huntley JS, Hall AC, Hall RH. Biochemical and physiological characteristics of HlyA, a pore-forming cytolysin of Vibrio cholerae serogroup O1. Toxicon : official journal of the International Society on Toxinology. 1997;35(4):515–27; doi: 10.1016/s0041-0101(96)00163-8.

81. Cinar HN, Kothary M, Datta AR, Tall BD, Sprando R, Bilecen K, et al. Vibrio cholerae hemolysin is required for lethality, developmental delay, and intestinal vacuolation in Caenorhabditis elegans. PLoS One. 2010;5(7):e11558; doi: 10.1371/journal.pone.0011558.

82. Chow KH, Ng TK, Yuen KY, Yam WC. Detection of RTX toxin gene in Vibrio cholerae by PCR. J Clin Microbiol. 2001;39(7):2594–7; doi: 10.1128/JCM.39.7.2594-2597.2001.

83. Halpern M, Gancz H, Broza M, Kashi Y. Vibrio cholerae hemagglutinin/protease degrades chironomid egg masses. Appl Environ Microbiol. 2003;69(7):4200–4; doi: 10.1128/AEM.69.7.4200-4204.2003.

84. Purdy AE, Balch D, Lizarraga-Partida ML, Islam MS, Martinez-Urtaza J, Huq A, et al. Diversity and distribution of cholix toxin, a novel ADP-ribosylating factor from Vibrio cholerae. Environ Microbiol Rep. 2010;2(1):198–207; doi: 10.1111/j.1758-2229.2010.00139.x.

85. Jorgensen R, Purdy AE, Fieldhouse RJ, Kimber MS, Bartlett DH, Merrill AR. Cholix toxin, a novel ADP-ribosylating factor from Vibrio cholerae. J Biol Chem. 2008;283(16):10671–8; doi: 10.1074/jbc.M710008200.

86. Nishibuchi M, Kaper JB. Thermostable direct hemolysin gene of Vibrio parahaemolyticus: a virulence gene acquired by a marine bacterium. Infect Immun. 1995;63(6):2093–9; doi: 10.1128/iai.63.6.2093-2099.1995.

87. Feldgarden M, Brover V, Gonzalez-Escalona N, Frye JG, Haendiges J, Haft DH, et al. AMRFinderPlus and the Reference Gene Catalog facilitate examination of the genomic links among antimicrobial resistance, stress response, and virulence. Sci Rep. 2021;11(1):12728; doi: 10.1038/s41598-021-91456-0.

88. Sebastian PJ, Schlesener C, Byrne BA, Miller M, Smith W, Batac F, et al. Antimicrobial resistance of Vibrio spp. from the coastal California system: discordance between genotypic and phenotypic patterns. Appl Environ Microbiol. 2025;91(3):e0180824; doi: 10.1128/aem.01808-24.

89. Hakonsholm F, Lunestad BT, Aguirre Sanchez JR, Martinez-Urtaza J, Marathe NP, Svanevik CS. Vibrios from the Norwegian marine environment: Characterization of associated antibiotic resistance and virulence genes. Microbiologyopen. 2020;9(9):e1093; doi: 10.1002/mbo3.1093.

90. Vandeputte M, Coppens S, Bossier P, Vereecke N, Vanrompay D. Genomic mining of Vibrio parahaemolyticus highlights prevalence of antimicrobial resistance genes and new genetic markers associated with AHPND and tdh + /trh + genotypes. BMC Genomics. 2024;25(1):178; doi: 10.1186/s12864-024-10093-9.

91. Baker-Austin C, Trinanes JA, Taylor NGH, Hartnell R, Siitonen A, Martinez-Urtaza J. Emerging Vibrio risk at high latitudes in response to ocean warming. Nat Clim Change. 2013;3(1):73–7; doi: 10.1038/Nclimate1628.

92. Witherall L, Wagley S, Butler C, Tyler CR, Temperton B. Genome Sequences of Four Vibrio parahaemolyticus Strains Isolated from the English Channel and the River Thames. Microbiol Resour Announc. 2019;8(24); doi: 10.1128/MRA.00392-19.

93. Ford CL, Powell A, Lau DYL, Turner AD, Dhanji-Rapkova M, Martinez-Urtaza J, et al. Isolation and characterization of potentially pathogenic Vibrio species in a temperate, higher latitude hotspot. Environ Microbiol Rep. 2020;12(4):424–34; doi: 10.1111/1758-2229.12858.

94. Hooban B, Whelan SO, Burke A, Lucey M, Tumeo A, Mulrooney C, et al. Emerging extraintestinal Vibrio infections in Ireland from clinical and marine sources, 2020-2022. Clin Microbiol Infect. 2026; doi: 10.1016/j.cmi.2026.02.013.

95. Ryder D, Adaway M, Batista FM, Wagley S, Powell A. The complete genome of Vibrio diabolicus isolated from coastal waters and Pacific oysters in England. Microbiol Resour Announc. 2025;14(7):e0131824; doi: 10.1128/mra.01318-24.

96. Investigations USfM. Gastroenteritis. UK Health Security Agency. 2026.

97. Hooban B, Whelan SO, Burke A, Lucey M, Tumeo A, Mulrooney C, et al. Emerging extraintestinal Vibrio infections in Ireland from clinical and marine sources, 2020-2022. Clin Microbiol Infect. 2026;32(7):1183–6; doi: 10.1016/j.cmi.2026.02.013.

98. Yang Y-Y, Lusk MG. Nutrients in Urban Stormwater Runoff: Current State of the Science and Potential Mitigation Options. Current Pollution Reports. 2018;4(2):112–27; doi: 10.1007/s40726-018-0087-7.

99. Conrad JW, Harwood VJ. Sewage Promotes Vibrio vulnificus Growth and Alters Gene Transcription in Vibrio vulnificus CMCP6. Microbiology spectrum. 2022;10(1):e0191321; doi: 10.1128/spectrum.01913-21.

100. Epa U: Nutrient Control Design Manual: state of technology review report. In.: EPA/600/R-09/012 EPA/600/R-09/012. US Environmental Protection Agency; 2009.

101. Singh V, Singh N, Rai SN, Kumar A, Singh AK, Singh MP, et al. Heavy Metal Contamination in the Aquatic Ecosystem: Toxicity and Its Remediation Using Eco-Friendly Approaches. Toxics. 2023;11(2); doi: 10.3390/toxics11020147.

102. Correa Velez KE, Norman RS. Transcriptomic Analysis Reveals That Municipal Wastewater Effluent Enhances Vibrio vulnificus Growth and Virulence Potential. Front Microbiol. 2021;12:754683; doi: 10.3389/fmicb.2021.754683.

103. Huijbers PM, Blaak H, de Jong MC, Graat EA, Vandenbroucke-Grauls CM, de Roda Husman AM. Role of the Environment in the Transmission of Antimicrobial Resistance to Humans: A Review. Environ Sci Technol. 2015;49(20):11993–2004; doi: 10.1021/acs.est.5b02566.

104. Goodwin KD, McNay M, Cao Y, Ebentier D, Madison M, Griffith JF. A multi-beach study of Staphylococcus aureus, MRSA, and enterococci in seawater and beach sand. Water Res. 2012;46(13):4195–207; doi: 10.1016/j.watres.2012.04.001.

105. Levin-Edens E, Soge OO, No D, Stiffarm A, Meschke JS, Roberts MC. Methicillin-resistant Staphylococcus aureus from Northwest marine and freshwater recreational beaches. FEMS Microbiol Ecol. 2012;79(2):412–20; doi: 10.1111/j.1574-6941.2011.01229.x.

106. Plano LR, Shibata T, Garza AC, Kish J, Fleisher JM, Sinigalliano CD, et al. Human-associated methicillin-resistant Staphylococcus aureus from a subtropical recreational marine beach. Microb Ecol. 2013;65(4):1039–51; doi: 10.1007/s00248-013-0216-1.

107. Schwartz T, Kohnen W, Jansen B, Obst U. Detection of antibiotic-resistant bacteria and their resistance genes in wastewater, surface water, and drinking water biofilms. FEMS Microbiol Ecol. 2003;43(3):325–35; doi: 10.1111/j.1574-6941.2003.tb01073.x.

108. Shuval H. Estimating the global burden of thalassogenic diseases: human infectious diseases caused by wastewater pollution of the marine environment. J Water Health. 2003;1(2):53–64.

109. Baker-Austin C, McArthur JV, Tuckfield RC, Najarro M, Lindell AH, Gooch J, et al. Antibiotic resistance in the shellfish pathogen Vibrio parahaemolyticus isolated from the coastal water and sediment of Georgia and South Carolina, USA. J Food Prot. 2008;71(12):2552–8.

110. Henderson JC, Herrera CM, Trent MS. AlmG, responsible for polymyxin resistance in pandemic Vibrio cholerae, is a glycyltransferase distantly related to lipid A late acyltransferases. J Biol Chem. 2017;292(51):21205–15; doi: 10.1074/jbc.RA117.000131.

111. Jordan A, Hill R, Turner A, Roberts T, Comber s: Assessing Options for Remediation of Contaminated Mine Site Drainage Entering the River Teign, Southwest England. In: Preprints. Preprints; 2020.

112. Simons B, Pirrie D, Rollinson GK, Shail R. Geochemical and mineralogical record of the impact of mining on the Teign Estuary, Devon, UK. 2025.

113. Zhang Y, Gu AZ, Cen T, Li X, He M, Li D, et al. Sub-inhibitory concentrations of heavy metals facilitate the horizontal transfer of plasmid-mediated antibiotic resistance genes in water environment. Environ Pollut. 2018;237:74–82; doi: 10.1016/j.envpol.2018.01.032.

114. Baker-Austin C, Wright MS, Stepanauskas R, McArthur JV. Co-selection of antibiotic and metal resistance. Trends Microbiol. 2006;14(4):176–82; doi: 10.1016/j.tim.2006.02.006.

115. Hill WE, Keasler SP, Trucksess MW, Feng P, Kaysner CA, Lampel KA. Polymerase chain reaction identification of Vibrio vulnificus in artificially contaminated oysters. Appl Environ Microbiol. 1991;57(3):707–11.

116. Rasko Leinonen HS, Martin Shumway, on behalf of the International Nucleotide Sequence Database Collaboration. The Sequence Read Archive. Nucleic Acids Res. 2011;39:D19–D21; doi: 10.1093/nar/gkq1019.

117. Felix Krueger FJ, Phil Ewels, Ebrahim Afyounian, Michael Weinstein, Benjamin Schuster-Boeckler, Gert Hulselmans, & sclamons.. FelixKrueger/TrimGalore: v0.6.10 - add default decompression path (0.6.10). Zenodo. 2023; doi: 10.5281/zenodo.7598955.

118. https://github.com/rrwick/Filtlong.

119. Wick RR, Judd LM, Gorrie CL, Holt KE. Unicycler: Resolving bacterial genome assemblies from short and long sequencing reads. PLoS Comput Biol. 2017;13(6):e1005595; doi: 10.1371/journal.pcbi.1005595.

120. Bouras G, Houtak G, Wick RR, Mallawaarachchi V, Roach MJ, Papudeshi B, et al. Hybracter: enabling scalable, automated, complete and accurate bacterial genome assemblies. Microb Genom. 2024;10(5); doi: 10.1099/mgen.0.001244.

121. Gurevich A, Saveliev V, Vyahhi N, Tesler G. QUAST: quality assessment tool for genome assemblies. Bioinformatics. 2013;29(8):1072–5; doi: 10.1093/bioinformatics/btt086.

122. Parks DH, Imelfort M, Skennerton CT, Hugenholtz P, Tyson GW. CheckM: assessing the quality of microbial genomes recovered from isolates, single cells, and metagenomes. Genome Res. 2015;25(7):1043–55; doi: 10.1101/gr.186072.114.

123. Benson DA, Karsch-Mizrachi I, Lipman DJ, Ostell J, Wheeler DL. GenBank. Nucleic Acids Res. 2005;33(Database issue):D34–8; doi: 10.1093/nar/gki063.

124. Sayers EW, Bolton EE, Brister JR, Canese K, Chan J, Comeau DC, et al. Database resources of the national center for biotechnology information. Nucleic Acids Res. 2022;50(D1):D20–D6; doi: 10.1093/nar/gkab1112.

125. Tatusova T, DiCuccio M, Badretdin A, Chetvernin V, Nawrocki EP, Zaslavsky L, et al. NCBI prokaryotic genome annotation pipeline. Nucleic Acids Res. 2016;44(14):6614–24; doi: 10.1093/nar/gkw569.

126. Shaw J, Yu YW. Rapid species-level metagenome profiling and containment estimation with sylph. Nat Biotechnol. 2025;43(8):1348–59; doi: 10.1038/s41587-024-02412-y.

127. Hernandez-Salmeron JE, Moreno-Hagelsieb G. FastANI, Mash and Dashing equally differentiate between Klebsiella species. PeerJ. 2022;10:e13784; doi: 10.7717/peerj.13784.

128. Jolley KA, Bray JE, Maiden MCJ. Open-access bacterial population genomics: BIGSdb software, the PubMLST.org website and their applications. Wellcome Open Res. 2018;3:124; doi: 10.12688/wellcomeopenres.14826.1.

129. T S. https://github.com/tseemann/mlst. 2026.

130. Shakya M, Ahmed SA, Davenport KW, Flynn MC, Lo CC, Chain PSG. Standardized phylogenetic and molecular evolutionary analysis applied to species across the microbial tree of life. Sci Rep. 2020;10(1):1723; doi: 10.1038/s41598-020-58356-1.

131. Price MN, Dehal PS, Arkin AP. FastTree 2--approximately maximum-likelihood trees for large alignments. PLoS One. 2010;5(3):e9490; doi: 10.1371/journal.pone.0009490.

132. O’Leary NA, Cox E, Holmes JB, Anderson WR, Falk R, Hem V, et al. Exploring and retrieving sequence and metadata for species across the tree of life with NCBI Datasets. Sci Data. 2024;11(1):732; doi: 10.1038/s41597-024-03571-y.

133. Katz LS, Griswold T, Morrison SS, Caravas JA, Zhang S, den Bakker HC, et al. Mashtree: a rapid comparison of whole genome sequence files. J Open Source Softw. 2019;4(44); doi: 10.21105/joss.01762.

134. Ondov BD, Treangen TJ, Melsted P, Mallonee AB, Bergman NH, Koren S, et al. Mash: fast genome and metagenome distance estimation using MinHash. Genome Biol. 2016;17(1):132; doi: 10.1186/s13059-016-0997-x.

135. Argimon S, Abudahab K, Goater RJE, Fedosejev A, Bhai J, Glasner C, et al. Microreact: visualizing and sharing data for genomic epidemiology and phylogeography. Microb Genom. 2016;2(11):e000093; doi: 10.1099/mgen.0.000093.

136. Team RC. R: A Language and Environment for Statistical Computing. Vienna: R Core Team. 2018. 2015.

137. Chen L, Yang J, Yu J, Yao Z, Sun L, Shen Y, et al. VFDB: a reference database for bacterial virulence factors. Nucleic Acids Res. 2005;33(Database issue):D325-8; doi: 10.1093/nar/gki008.

138. Seemann T. n.d. Abricate (version 0.8). Github. Accessed March 7, 2021. https://githubcom/tseemann/abricate. 2021.

139. Oliver JD, Colwell RR. Extractable lipids of gram-negative marine bacteria: phospholipid composition. J Bacteriol. 1973;114(3):897–908; doi: 10.1128/jb.114.3.897-908.1973.

140. Sheikh HI, Najiah M, Fadhlina A, Laith AA, Nor MM, Jalal KCA, et al. Temperature Upshift Mostly but not Always Enhances the Growth of Vibrio Species: A Systematic Review. Frontiers in Marine Science. 2022;Volume 9 - 2022; doi: 10.3389/fmars.2022.959830.

141. Sprouffske K, Wagner A. Growthcurver: an R package for obtaining interpretable metrics from microbial growth curves. BMC Bioinformatics. 2016;17:172; doi: 10.1186/s12859-016-1016-7.

142. Bier N, Schwartz K, Guerra B, Strauch E. Survey on antimicrobial resistance patterns in Vibrio vulnificus and Vibrio cholerae non-O1/non-O139 in Germany reveals carbapenemase-producing Vibrio cholerae in coastal waters. Front Microbiol. 2015;6:1179; doi: 10.3389/fmicb.2015.01179.

143. Karatuna O, Matuschek E, Ahman J, Caidi H, Kahlmeter G. Vibrio species: development of EUCAST susceptibility testing methods and MIC and zone diameter distributions on which to determine clinical breakpoints. J Antimicrob Chemother. 2024;79(2):375–82; doi: 10.1093/jac/dkad391.

