## Supplementary Table 1 and 2 for "Environmental Drivers and Distribution of Pathogenic *Vibrio* Species in the Teign Estuary, UK"

**Supplementary Table S1:** Water, Sediment, Shellfish and Seaweed samples tested in this study between 2023 and 2025.

^A^ : type of sample tested: A – Water, B – Sediment, C – *Cerastoderma edule* (common cockle), D – *Magallana gigas* (Pacific oyster), E - *Spisula solida* (surf Clam), F – *Mytilus edulis* (blue mussel), G –Seaweed

| Sample number | Date of Isolation | Type of sample ^A^ | Total no of Vibrios detected CFU/ml or CFU/g | *Vibrio* species isolated |
| --- | --- | --- | --- | --- |
| EXE 07/2023 | Sept 2023 | A | 20 | ***V. diabolicus*** |
| EXE 08/2023 | Sept 2023 | B | 3433 | ***V. diabolicus*** |
| EXE 09/2023 | Sept 2023 | C | 3200 | ***V. diabolicus*** |
| EXE 10/2023 | Sept 2023 | D | 8500 | ***V. diabolicus*** |
| EXE 01/2024 | May 2024 | A | <20 | ***V. alginolyticus*** |
| EXE 02/2024 | May 2024 | B | 150 | *V. alginolyticus* |
| EXE 03/2024 | May 2024 | C | 200 | *V. alginolyticus* |
| EXE 04/2024 | May 2024 | D | 1150 | *Vibrio* species detected |
| EXE 05/2024 | June 2024 | A | <20 | *V. alginolyticus, V. cholerae* |
| EXE 06/2024 | June 2024 | B | 50 | ***V. diabolicus*** |
| EXE 07/2024 | June 2024 | E | 50 | *V. alginolyticus* |
| EXE 08/2024 | June 2024 | C | 200 | ***V. diabolicus*** |
| EXE 09/2024 | June 2024 | D | 300 | ***V. alginolyticus*** |
| EXE 10/2024 | July 2024 | A | <20 | *V. alginolyticus* |
| EXE 11/2024 | July 2024 | B | 600 | *V. alginolyticus* |
| EXE 14/2024 | August 2024 | A | 30 | *V. alginolyticus* |
| EXE 15/2024 | August 2024 | B | 750 | *V. alginolyticus,* ***V. parahaemolyticus*** |
| EXE 16/2024 | August 2024 | D | 8350 | *V. alginolyticus* |
| EXE 17/2024 | August 2024 | C | 12350 | *V. alginolyticus* |
| EXE 18/2024 | August 2024 | F | 2900 | *V. alginolyticus* |
| EXE 19/2024 | August 2024 | E | 2050 | ***V. diabolicus*** *V. parahaemolyticus* |
| EXE 20/2024 | August 2024 | G | 350 | ***V. diabolicus*** |
| EXE 21/2024 | Sept 2024 | A | 20 | *V. parahaemolyticus, V. alginolyticus* |
| EXE 22/2024 | Sept 2024 | B | 300 | ***V. jasicida,*** *V. parahaemolyticus, V. alginolyticus* |
| EXE 23/2024 | Sept 2024 | C | 700 | ***V. diabolicus****,* ***V. parahaemolyticus*** |
| EXE 24/2024 | Sept 2024 | F | 1850 | *V. alginolyticus,* |
| EXE 25/2024 | Sept 2024 | E | 500 | *V. alginolyticus* ***V. parahaemolyticus*** |
| EXE 26/2024 | Sept 2024 | G | 4100 | ***V. parahaemolyticus,*** *V. alginolyticus* |
| EXE 29/2024 | Oct 2024 | A | <20 | *V. alginolyticus* |
| EXE 30/2024 | Oct 2024 | B | <20 | ***V. alginolyticus,*** ***V. jasicida*** |
| EXE 31/2024 | Oct 2024 | C | 200 | *V. alginolyticus* |
| EXE 32/2024 | Oct 2024 | F | 50 | *V. alginolyticus,* ***V. parahaemolyticus,*** |
| EXE 33/2024 | Oct 2024 | E | 100 | ***V. diabolicus,*** ***V. parahaemolyticus*** |
| EXE 34/2024 | Oct 2024 | G | <20 | ***V. diabolicus,*** ***V. parahaemolyticus,*** |
| EXE 35/2024 | Nov 2024 | A | 0 |  |
| EXE 36/2024 | Nov 2024 | B | 0 |  |
| EXE 37/2024 | Nov 2024 | G | 0 |  |
| EXE 38/2024 | Nov 2024 | E | 0 |  |
| EXE 39/2024 | Nov 2024 | F | 0 |  |
| EXE 40/2024 | Nov 2024 | C | 0 |  |
| EXE 41/2024 | Nov 2024 | D | 0 |  |
| EXE 42/2024 | Dec 2024 | A | 0 |  |
| EXE 43/2024 | Dec 2024 | B | 0 |  |
| EXE 44/2024 | Dec 2024 | C | 0 |  |
| EXE 45/2024 | Dec 2024 | E | <20 | ***V. cholerae*** |
| EXE 01/2025 | Jan 2025 | A | 10 | ***V. anguillarum*** |
| EXE 02/2025 | Jan 2025 | B | 0 |  |
| EXE 03/2025 | Jan 2025 | C | 0 | *Vibrio species, V. alginolyticus,* |
| EXE 04/2025 | Jan 2025 | G | 0 |  |
| EXE 05/2025 | Feb 2025 | A | 0 |  |
| EXE 06/2025 | Feb 2025 | B | 0 |  |
| EXE 07/2025 | Feb 2025 | C | 0 |  |
| EXE 08/2025 | March 2025 | A | 0 |  |
| EXE 09/2025 | March 2025 | B | 0 |  |
| EXE 10/2025 | March 2025 | C | 0 |  |
| EXE 11/2025 | April 2025 | A | 0 |  |
| EXE 12/2025 | April 2025 | B | <20 | ***V. diabolicus,*** *V. aestuarianus* |
| EXE 13/2025 | April 2025 | C | <20 | ***V. aestuarianus,*** ***V. jasicida*** |
| EXE 14/2025 | May 2025 | A | 0 |  |
| EXE 15/2025 | May 2025 | B | 0 |  |
| EXE 16/2025 | May 2025 | C | <20 | *V. alginolyticus* |
| EXE 17/2025 | May 2025 | E | <20 | *V. alginolyticus* |
| EXE 20/2025 | June 2025 | A | <20 | ***V. alginolyticus*** |
| EXE 21/2025 | June 2025 | B | <20 | ***V. alginolyticus*** |
| EXE 22/2025 | June 2025 | C | 250 | ***V. diabolicus,*** ***V. jasicida*** |
| EXE 23/2025 | June 2025 | E | 50 | ***Vibrio species****, V. alginolyticus* |
| EXE 24/2025 | June 2025 | F | <20 | ***V. alginolyticus,*** ***V. parahaemolyticus*** |
| EXE 25/2025 | June 2025 | D | 1550 | ***V. diabolicus*** |
| EXE 26/2025 | July 2025 | A | <20 | ***V. jasicida,*** ***V. neptunius*** |
| EXE 27/2025 | July 2025 | B | 150 | ***V. jasicida*,** ***V. diabolicus*** |
| EXE 28/2025 | July 2025 | C | 3100 | *V. alginolyticus,* ***V. alfacsensis****,* ***V. pacinii*** |
| EXE 29/2025 | July 2025 | E | 3200 | ***V. jasicida,*** ***V. diabolicus****,* ***V. parahaemolyticus*** |
| EXE 30/2025 | July 2025 | F | 550 | ***V. alginolyticus,*** ***V. parahaemolyticus*** |
| EXE 31/2025 | August 2025 | A | <20 | *V. alginolyticus* |
| EXE 32/2025 | August 2025 | B | 400 | *V. alginolyticus, V. parahaemolyticus* |
| EXE 33/2025 | August 2025 | C | 2850 | *V. alginolyticus* |
| EXE 34/2025 | August 2025 | E | 350 | *V. alginolyticus, V. parahaemolyticus* |
| EXE 35/2025 | August 2025 | G | 0 |  |
| EXE 37/2025 | Sept 2025 | A | <20 | *V. alginolyticus* |
| EXE 38/2025 | Sept 2025 | B | 200 | *V. alginolyticus* |
| EXE 39/2025 | Sept 2025 | D | 8400 | *V. alginolyticus, V. parahaemolyticus* |
| EXE 40/2025 | Sept 2025 | G | 0 |  |
| EXE 41/2025 | Sept 2025 | G | 200 | *V. alginolyticus* |
| EXE 42/2025 | Sept 2025 | A | <20 | *V. alginolyticus* |
| EXE 43/2025 | Oct 2025 | A | <20 | *V. alginolyticus* |
| EXE 44/2025 | Oct 2025 | B | 0 |  |
| EXE 45/2025 | Oct 2025 | C | >20 | *V. alginolyticus, V. parahaemolyticus* |
| EXE 46/2025 | Oct 2025 | D | >20 | *V. parahaemolyticus* |
| EXE 47/2025 | Oct 2025 | E | >20 | *V. parahaemolyticus* |
| EXE 48/2025 | Oct 2025 | F | 0 |  |
| EXE 49/2025 | Oct 2025 | G | 0 |  |

**Supplementary Table S2: Dose range injected to *G. mellonella***

| **Strain** | **Species** | **Dose Range** |
| --- | --- | --- |
| EXE 09/2024-B | *V. alginolyticus* | 141.5-210 |
| EXE 30/2025 -L | *V. alginolyticus* | 100-965 |
| Guernsey strain | *V. alginolyticus* | 200-430 |
| EXE 07/2023-A4 | *V. diabolicus* | 73-1935 |
| EXE 20/2024-J | *V. diabolicus* | 73-915 |
| EXE 27/2025 - D | *V. diabolicus* | 85-835 |
| EXE 05/2024-A | *V. cholerae* | 83-330 |
| EXE 45/2024-E | *V. cholerae* | 140-270 |
| EXE 19/2024-C | *V. parahaemolyticus* | 141.5-600 |
| EXE 24/2025 - E | *V. parahaemolyticus* | 80-1435 |
| G35 Clinical | *V. parahaemolyticus* | 93-765 |
