## Supplementary Table 3 for "Environmental Drivers and Distribution of Pathogenic *Vibrio* Species in the Teign Estuary, UK"

**Supplementary Table S3: Antibiotic susceptibilities.** The table shows the results of MIC assays for 13 Vibrio strains isolated from the Teign Estuary and one clinical isolated from a wound infection for Guernsey. AMP: ampicillin, Kan: kanamycin, Cm: chloramphenicol, Tp: trimethoprim, Cf: ceftazidime, Cp: ciprofloxacin, Dox: doxycycline, Gen: gentamicin.

| Strain | Date of Isolation | Species | Antibiotic MIC | | | | | | | | | MAR Index (>0.2) |
| --- | --- | --- | --- | --- | --- | --- | --- | --- | --- | --- | --- | --- |
|  |  |  | **AMP** | **Kan** | **Cm** | **Tp** | **Cf** | **Cp** | **Dox** | **Gent** |  | |
| Guernsey strain | 2011 | *V. alginolyticus* |  |  |  |  |  |  |  |  | 0.625 | |
| EXE 07-2023 A4 | 2023 | *V. diabolicus* | 1024 | 16 | 32 | 256 | 1024 | 64 | >1 | 8 | 0.75 | |
| EXE 09/2024-B | 2024 | *V. alginolyticus* | 1024 | 16 | 32 | 1024 | 1024 | 4 | >1 | 16 | 0.75 | |
| EXE 01/2024-7 | 2024 | *V. alginolyticus* | 1024 | 16 | 4 | 1024 | 1024 | 4 | >1 | 16 | 0.625 | |
| EXE 23/2024-K | 2024 | *V. diabolicus* | 1024 | 16 | 8 | 512 | 1024 | 4 | >1 | 8 | 0.625 | |
| EXE 06/2024-E | 2024 | *V. diabolicus* | 1024 | 16 | 8 | 1024 | 1024 | 4 | >1 | 8 | 0.625 | |
| EXE 15/2024-J | 2024 | *V. parahaemolyticus* | 1024 | 16 | 64 | 1024 | 1024 | 4 | >1 | 16 | 0.75 | |
| EXE 19/2024-C | 2024 | *V. parahaemolyticus* | 1024 | 16 | 128 | 1024 | 1024 | 4 | >1 | 16 | 0.75 | |
| EXE 25/2024-H | 2024 | *V. parahaemolyticus* | 1024 | 16 | 256 | 1024 | 1024 | 4 | >1 | 8 | 0.75 | |
| EXE 33/2024-F | 2024 | *V. parahaemolyticus* | 1024 | 16 | 8 | 1024 | 1024 | 4 | >1 | 16 | 0.625 | |
| EXE 20/2025 - A | 2025 | *V. alginolyticus* | 1024 | 16 | >1 | 2 | 4 | >1 | >1 | 8 | 0.5 | |
| EXE 20/2025 - E | 2025 | *V. parahaemolyticus* | 1024 | 32 | >1 | 256 | 4 | >1 | >1 | 8 | 0.625 | |
| EXE 29/2025 -HH | 2025 | *V. parahaemolyticus* | 1024 | 32 | >1 | 1024 | 1024 | >1 | >1 | 8 | 0.625 | |
| EXE 13/2025-H | 2025 | *V. aestuarianus* | 1024 | 32 | >1 | >1 | 1024 | >1 | >1 | 8 | 0.5 | |
