## Supplementary Table 4 for "Environmental Drivers and Distribution of Pathogenic *Vibrio* Species in the Teign Estuary, UK"

**Supplementary Table S4. Genome sequencing for each of the *Vibrio* isolates sequenced in this study.**

| Run | Bases | Instrument | isolate | Organism |
| --- | --- | --- | --- | --- |
| SRR38114694 | 281,355,548 | PromethION | EXE23 2025 L | *Vibrio* sp. |
| SRR38114727 | 1,417,424,048 | PromethION | EXE13 2025 H | *Vibrio aestuarianus* |
| SRR38114716 | 2,334,101,174 | PromethION | EXE28 2025 A | *Vibrio alfacsensis* |
| SRR38114675 | 2,208,076,285 | PromethION | EXE01 2024 7 | *Vibrio alginolyticus* |
| SRR38114729 | 1,194,078,450 | PromethION | EXE09 2024 B | *Vibrio alginolyticus* |
| SRR38114695 | 298,597,469 | PromethION | EXE20 2025 B | *Vibrio alginolyticus* |
| SRR38114723 | 6,376,501,672 | PromethION | EXE21 2025 F | *Vibrio alginolyticus* |
| SRR38158597 | 16,532,419 | PromethION | EXE24 2025 A | *Vibrio alginolyticus* |
| SRR38114712 | 1,392,527,740 | PromethION | EXE30 2024 F | *Vibrio alginolyticus* |
| SRR38114710 | 797,585,428 | PromethION | EXE30 2025 L | *Vibrio alginolyticus* |
| SRR38114677 | 616,848,300 | Illumina NovaSeq X | Guernsey | *Vibrio alginolyticus* |
| SRR38114678 | 766,613,082 | PromethION | Guernsey | *Vibrio alginolyticus* |
| SRR38114701 | 289,333,669 | PromethION | EXE01 2025 M | *Vibrio anguillarum* |
| SRR38114674 | 99,753,168 | PromethION | EXE05 2024 A | *Vibrio cholerae* |
| SRR38114687 | 607,120,500 | Illumina NovaSeq X | EXE45 2024 E | *Vibrio cholerae* |
| SRR38114679 | 643,351,652 | PromethION | EXE45 2024 E | *Vibrio cholerae* |
| SRR38114730 | 2,729,391,809 | PromethION | EXE06 2024 E | *Vibrio diabolicus* |
| SRR38114690 | 1,818,297,000 | Illumina NovaSeq 6000 | EXE07 2023 A1 | *Vibrio diabolicus* |
| SRR38114689 | 1,699,808,100 | Illumina NovaSeq 6000 | EXE07 2023 A4 | *Vibrio diabolicus* |
| SRR38114688 | 2,024,748,600 | Illumina NovaSeq 6000 | EXE08 2023 B5 | *Vibrio diabolicus* |
| SRR38114699 | 386,182,255 | PromethION | EXE08 2024 A | *Vibrio diabolicus* |
| SRR38114686 | 2,282,127,600 | Illumina NovaSeq 6000 | EXE09 2023 D7 | *Vibrio diabolicus* |
| SRR38114685 | 2,010,215,400 | Illumina NovaSeq 6000 | EXE10 2023 G3 | *Vibrio diabolicus* |
| SRR38114728 | 2,857,892,034 | PromethION | EXE12 2025 A | *Vibrio diabolicus* |
| SRR38114725 | 757,573,890 | PromethION | EXE19 2024 M | *Vibrio diabolicus* |
| SRR38114724 | 543,480,040 | PromethION | EXE20 2024 J | *Vibrio diabolicus* |
| SRR38114721 | 3,424,675,450 | PromethION | EXE22 2025 T1A | *Vibrio diabolicus* |
| SRR38114676 | 732,087,538 | PromethION | EXE23 2024 K | *Vibrio diabolicus* |
| SRR38158595 | 2,410,568 | PromethION | EXE25 2025 25N | *Vibrio diabolicus* |
| SRR38114717 | 2,535,377,060 | PromethION | EXE27 2025 D | *Vibrio diabolicus* |
| SRR38114713 | 1,038,399,716 | PromethION | EXE29 2025 V | *Vibrio diabolicus* |
| SRR38114692 | 653,564,386 | PromethION | EXE32 2024 G | *Vibrio diabolicus* |
| SRR38114706 | 145,412,191 | PromethION | EXE33 2024 C | *Vibrio diabolicus* |
| SRR38114691 | 501,633,499 | PromethION | EXE33 2024 E | *Vibrio diabolicus* |
| SRR38114704 | 281,511,229 | PromethION | EXE34 2024 C | *Vibrio diabolicus* |
| SRR38114700 | 617,222,231 | PromethION | EXE03 2025 A | *Vibrio jasicida* |
| SRR38114697 | 503,634,921 | PromethION | EXE13 2025 A | *Vibrio jasicida* |
| SRR38114726 | 5,189,869,802 | PromethION | EXE13 2025 P | *Vibrio jasicida* |
| SRR38114683 | 301,306,323 | PromethION | EXE22 2024 B | *Vibrio jasicida* |
| SRR38114731 | 601,080,900 | Illumina NovaSeq X | EXE22 2024 B | *Vibrio jasicida* |
| SRR38158598 | 280,905,447 | PromethION | EXE22 2025 DG1+LF | *Vibrio jasicida* |
| SRR38114722 | 306,717,939 | PromethION | EXE22 2025 DG3A | *Vibrio jasicida* |
| SRR38114702 | 634,858,641 | PromethION | EXE22 2025 IVF5+LA | *Vibrio jasicida* |
| SRR38114719 | 396,189,996 | PromethION | EXE26 2025 A | *Vibrio jasicida* |
| SRR38114718 | 499,464,283 | PromethION | EXE27 2025 A | *Vibrio jasicida* |
| SRR38114714 | 1,962,367,369 | PromethION | EXE29 2025 K | *Vibrio jasicida* |
| SRR38114711 | 748,022,378 | PromethION | EXE30 2024 G | *Vibrio jasicida* |
| SRR38158594 | 547,067,820 | PromethION | EXE26 2025 C | *Vibrio neptunius* |
| SRR38158593 | 918,722,736 | PromethION | EXE28 2025 W | *Vibrio pacinii* |
| SRR38114696 | 574,187,700 | Illumina NovaSeq X | EXE15 2024 J | *Vibrio parahaemolyticus* |
| SRR38114684 | 909,847,933 | PromethION | EXE19 2024 C | *Vibrio parahaemolyticus* |
| SRR38114732 | 2,855,509,800 | Illumina NovaSeq X | EXE19 2024 C | *Vibrio parahaemolyticus* |
| SRR38114682 | 526,302,865 | PromethION | EXE23 2024 1 | *Vibrio parahaemolyticus* |
| SRR38114720 | 719,169,600 | Illumina NovaSeq X | EXE23 2024 1 | *Vibrio parahaemolyticus* |
| SRR38158596 | 6,168,598 | PromethION | EXE24 2025 E | *Vibrio parahaemolyticus* |
| SRR38114709 | 634,501,800 | Illumina NovaSeq X | EXE25 2024 H | *Vibrio parahaemolyticus* |
| SRR38114681 | 652,688,133 | PromethION | EXE25 2024 H | *Vibrio parahaemolyticus* |
| SRR38114698 | 600,890,700 | Illumina NovaSeq X | EXE26 2024 C | *Vibrio parahaemolyticus* |
| SRR38114680 | 925,627,375 | PromethION | EXE26 2024 C | *Vibrio parahaemolyticus* |
| SRR38114715 | 1,609,639,321 | PromethION | EXE29 2025 HH | *Vibrio parahaemolyticus* |
| SRR38114693 | 258,125,941 | PromethION | EXE30 2025 R | *Vibrio parahaemolyticus* |
| SRR38114708 | 1,215,031,592 | PromethION | EXE30 2025 S | *Vibrio parahaemolyticus* |
| SRR38114707 | 240,815,248 | PromethION | EXE32 2024 G | *Vibrio parahaemolyticus* |
| SRR38114705 | 1,487,276,743 | PromethION | EXE33 2024 F | *Vibrio parahaemolyticus* |
| SRR38114703 | 2,638,884,735 | PromethION | EXE34 2024 P | *Vibrio parahaemolyticus* |
