## Supplementary Table 15 for "Environmental Drivers and Distribution of Pathogenic *Vibrio* Species in the Teign Estuary, UK"

**Supplementary Table S5: PCR primers and conditions for identifying V. diabolicus isolates.**

| Gene |  | Sequence (5’-3’) | Annealing Temp. (°C) | Size  (bp) |
| --- | --- | --- | --- | --- |
| Hypothetical protein (ACY49842.1) | Forward | ATGCAGGTTGTAGCACGTGCCTAA | 55 | 225 |
|  | Reverse | TTATTGATGGTCCTGTCGGATTTTTATC |  |  |
| Ornithine cyclodeaminase (ACY53325.1) | Forward | TCTCAGATAGAGCAAGTGACTCAG | 55 | 960 |
|  | Reverse | TTACATCGTGGATTGAATGCTGGT |  |  |
| Hydroxyectoine utilisation dehydratase (ACY53324.1) | Forward | ATGGAAACACAAGCAATCTTTGCC | 54 | 975 |
|  | Reverse | TTATTCTTTGACTTGGCGGCCAAT |  |  |
| Phosphoesterase (ACY53763.1) | Forward | TCAAAAAGTGCAGTGGCAACTTTA | 53 | 2234 |
|  | Reverse | TATGACTGCAACCAAACAGGG |  |  |
